# The cell cycle in *Staphylococcus aureus* is regulated by an amidase that controls peptidoglycan synthesis

**DOI:** 10.1101/634089

**Authors:** Truc Do, Kaitlin Schaefer, Ace George Santiago, Kathryn A. Coe, Pedro B. Fernandes, Daniel Kahne, Mariana G. Pinho, Suzanne Walker

## Abstract

Bacteria are protected by a polymer of peptidoglycan that serves as an exoskeleton. In *Staphylococcus aureus*, the enzymes that assemble peptidoglycan move during the cell cycle from the periphery, where they are active during growth, to the division site where they build the partition between daughter cells. But how peptidoglycan synthesis is regulated throughout the cell cycle is not understood. Here we identify a membrane protein complex that spatially regulates *S. aureus* peptidoglycan synthesis. This complex consists of an amidase that removes peptide chains from uncrosslinked peptidoglycan and a partner protein that controls its activity. Typical amidases act after cell division to hydrolyze peptidoglycan between daughter cells so they can separate. However, we show that this amidase controls cell growth. In its absence, excess peptidoglycan synthesis occurs at the cell periphery, causing cells to grow so large that cell division is defective. We show that cell growth and division defects due to loss of this amidase can be mitigated by attenuating the polymerase activity of the major *S. aureus* peptidoglycan synthase. Our findings lead to a model wherein the amidase complex regulates the density of peptidoglycan assembly sites to control peptidoglycan synthase activity at a given cellular location. Removal of peptide chains from peptidoglycan at the cell periphery promotes synthase movement to midcell during cell division. This mechanism ensures that cell expansion is properly coordinated with cell division.

## Main text

The bacterial cell wall, which is largely composed of a polymer called peptidoglycan, is essential for survival. The chemical steps in cell wall biosynthesis are conserved, but how the activities of peptidoglycan biosynthetic enzymes are coordinated with growth and division is unclear^1,2^. To make peptidoglycan, glycan chains are first polymerized from a disaccharide-peptide precursor, Lipid II, and then crosslinked to the existing matrix via their attached peptides (Fig. 1a). Crosslinking is carried out by the transpeptidase domain of penicillin-binding proteins (PBPs) using a mechanism that involves formation of a covalent adduct between the enzymes and the stem peptides on newly synthesized peptidoglycan^3^. There are two modes of peptidoglycan synthesis during the cell cycle. One mode occurs at the cell periphery and leads to an expansion in cell size; the other occurs during cell division when a cross wall, or septum, is formed between two daughter cells^4–6^. The first protein that assembles at the division site is the highly conserved tubulin homolog FtsZ, which forms a ring-shaped structure called the Z-ring that serves as a scaffold for the ordered recruitment of the rest of the divisome^7^. In *Staphylococcus aureus*, proper placement of FtsZ depends on controlling cell size^8–10^, which in turn requires spatial control over peptidoglycan synthesis.

**Figure 1:**
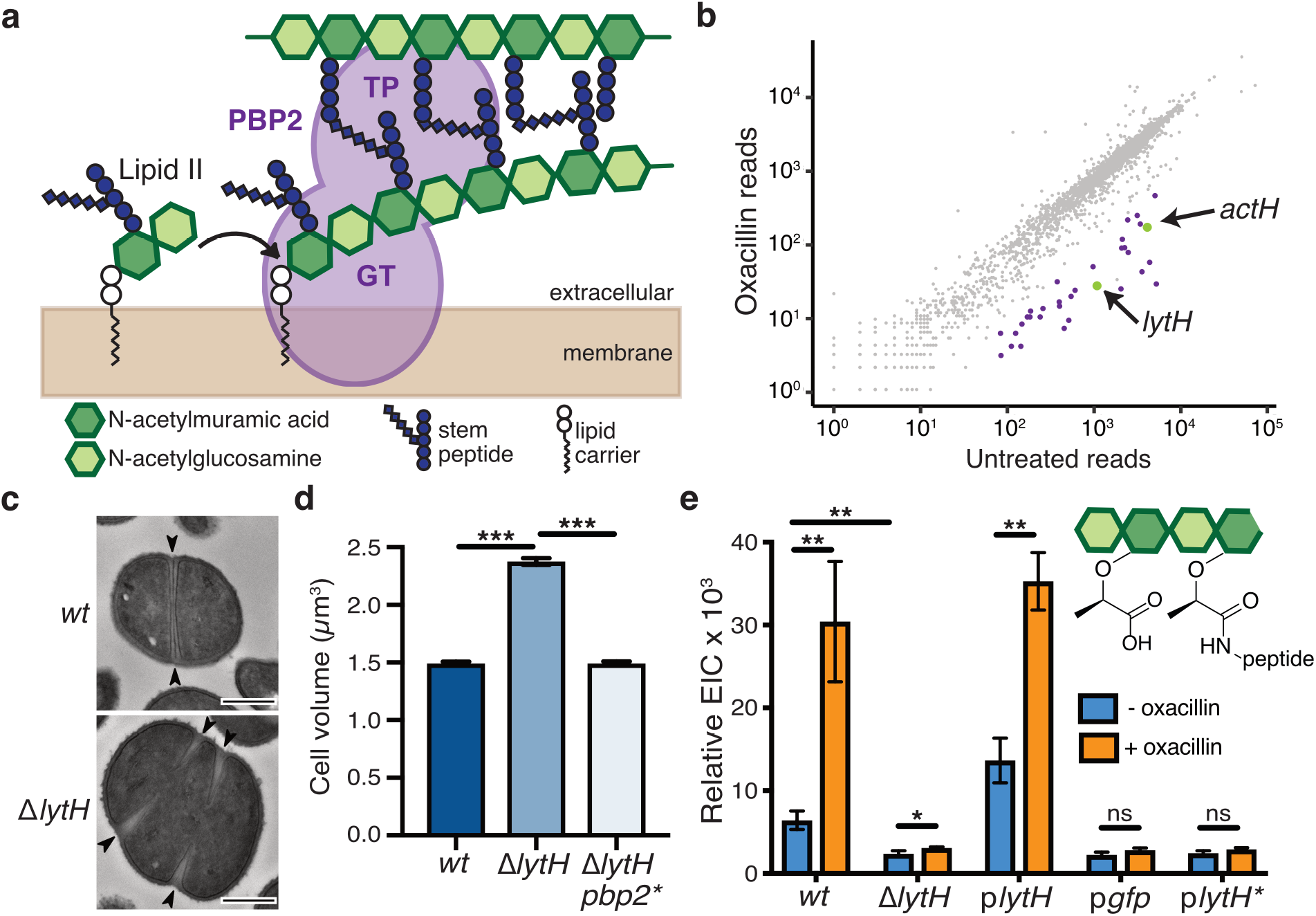
LytH is an amidase involved in cell division. **a**, Peptidoglycan is assembled from Lipid II. PBP2 catalyzes the polymerization (GT, glycosyltransferase domain) of Lipid II and crosslinking (TP, transpeptidase domain) of glycan strands into the existing matrix. **b**, Transposon sequencing reads in *lytH* and *actH* (green) were depleted with oxacillin. The plot shows transposon reads for each gene (dot) of strain HG003 under the two conditions. Significantly depleted genes are denoted in green and purple. **c**, Transmission electron micrographs: WT cells have cross walls at midcell (arrows), but Δ*lytH* cells have multiple partial septa and are larger. Scale bars, 500 nm. **d**, Cell size represented as mean ± SEM (~700 non-dividing cells/strain). *pbp2\** denotes *pbp2^F158L^*. P-values were determined by two-sided Mann-Whitney U tests. \*\*\**P* < 0.001. **e**, Sacculi analysis: A tetrasaccharide-monopeptide was more abundant in cells expressing *lytH* and increased under oxacillin treatment. p*lytH*, p*gfp*, and p*lytH\** denote strains expressing *lytH^WT^*, *gfp*, or *lytH^D195A^*, respectively, from a plasmid in a ∆*lytH* background. Data represent mean ± SD of the relative extracted ion count (EIC) from three experiments. P-values were determined by unpaired, two-tailed *t*-tests. \**P* < 0.05; \*\**P* < 0.01; ns, not significant.

Here, we used a transposon screen to identify possible regulators of peptidoglycan synthesis in *S. aureus*. We discovered a gene encoding a putative amidase, LytH, and showed that deletion of this gene caused dramatic phenotypes resulting from dysregulated coordination of cell growth with cell division. Reconstitution of LytH amidase activity required a membrane protein partner, ActH, that forms a stable complex with the amidase. Using uncrosslinked and crosslinked substrates made *in vitro*, we found that the purified protein complex only removes uncrosslinked stem peptides; cellular experiments were consistent with this preference. We used fluorescence microscopy to show that excess peptidoglycan synthesis occurs at the cell periphery when LytH is absent. Genetics combined with biochemistry showed that decreasing activity of the major peptidoglycan synthase, PBP2, corrects the cell size and division defects due to *lytH* deletion. We also showed that PBP2 accumulates at the cell periphery in the absence of LytH, which is likely responsible for the excess peptidoglycan synthesis there. Taken together, our findings lead to a model in which the LytH-ActH complex acts to remove free stem peptides from peptidoglycan in order to control the density of peptidoglycan assembly sites, and thereby to control cell expansion.

## Results

### A transposon screen identified *lytH* as important in cell size control and divisome placement

To identify new regulators of peptidoglycan synthesis in *S. aureus*, we performed a transposon screen^11^ using sublethal oxacillin to partially inhibit transpeptidase activity. We hypothesized that genes whose absence was synthetically lethal with partial loss of transpeptidase activity might regulate peptidoglycan synthesis. Reads due to transposon insertions were depleted in a number of genes, but we focused on *lytH*, which encodes a putative amidase, because inactivation of a cell wall hydrolase typically protects, rather than sensitizes, cells towards a β-lactam^12^ (Fig. 1b and Supplementary Table 1). A spot titer assay confirmed that *lytH* deletion sensitized cells to oxacillin (Supplementary Fig. 1). Amidases are hydrolytic enzymes that cleave stem peptides from peptidoglycan to separate daughter cells following cell division^13^. The Δ*lytH* mutant exhibited division defects, but they were not the cell separation defects previously observed for amidases. Instead, deletion of *lytH* led to misplacement of nascent septa (Fig. 1c) and larger cell size (Fig. 1d). The nature of the defects implied that LytH affects cell division at an earlier stage than amidases that effect cell separation.

### Muropeptide analysis confirmed that LytH has amidase activity

To assess LytH amidase activity, we compared the chemical composition of sacculi from wild-type (WT) and Δ*lytH* cells. Purified sacculi were digested with mutanolysin to generate muropeptides for LC-MS analysis (Supplementary Fig. 2). We then searched for muropeptides consistent with amidase processing and identified a species having an exact mass that suggested structure X^14^ (Supplementary Fig. 3a). This species was more abundant in sacculi from cells expressing a catalytically-active copy of *lytH* ^15^ than from Δ*lytH* cells (Fig. 1e). Using LC-MS/MS, we confirmed the identity of this species, which is a tetrasaccharide with only a single stem peptide attached (Supplementary Fig. 3b,c). We were able to detect this tetrasaccharide-monopeptide species because mutanolysin cannot cleave MurNAc-GlcNAc linkages unless the MurNAc residue carries a stem peptide^16^. The tetrasaccharide-monopeptide likely represents only a fraction of LytH products, as longer glycan chains extensively lacking stem peptides would not be detected with this method.

### LytH requires a partner protein for amidase activity

Having *in vivo* evidence of LytH amidase activity, we attempted to reconstitute its activity *in vitro*, but these efforts were unsuccessful (Supplementary Fig. 4). We speculated that LytH requires a partner protein for activation. To identify potential activators, we performed co-immunoprecipitation and isolated a polytopic membrane protein containing seven predicted transmembrane helices and a C-terminal domain with three tetratricopeptide repeats (TPRs) (Fig. 2a and Supplementary Fig. 5). This membrane protein was previously designated Rbd because it is a putative rhomboid protease^10^ (Supplementary Table 2). A mutant lacking this gene was sensitive to oxacillin (Fig. 1b and Supplementary Fig. 6) and displayed cell division defects similar to the ∆*lytH* mutant (Fig. 2b), supporting a role for the uncharacterized protein in activating LytH.

**Figure 2:**
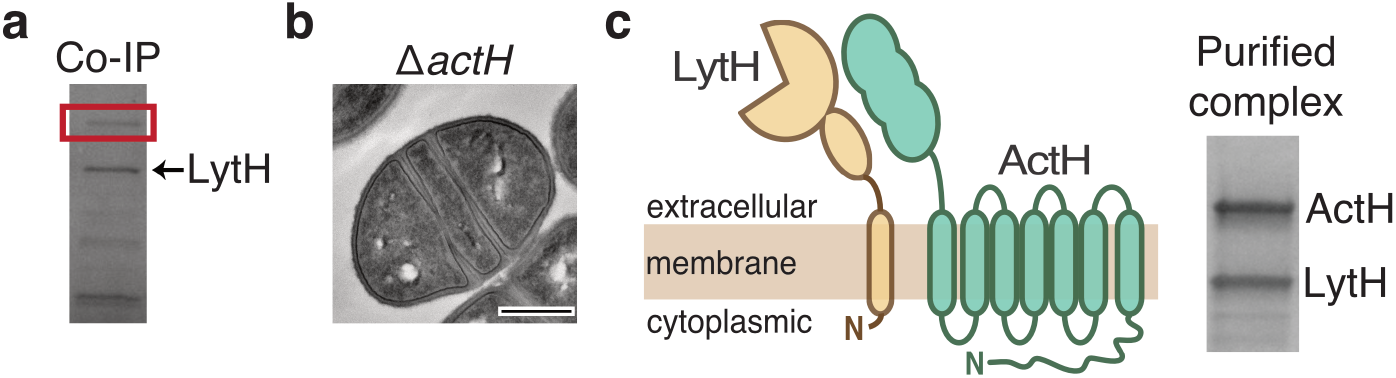
LytH forms a stable membrane complex with a partner protein. **a**, A pull-down identified ActH (red box) as a LytH-interaction partner. **b**, An *actH* deletion mutant exhibited septal defects similar to ∆*lytH* division defects. Scale bar, 500 nm. **c**, LytH and ActH purified as a complex after co-expression in *E. coli*. A schematic of the predicted topology is shown.

We co-expressed LytH and its putative protein partner and purified a stable complex (Fig. 2c and Supplementary Fig. 7a). We first tested amidase activity of the purified complex on peptidoglycan polymers prepared from native *S. aureus* Lipid II^17^ using a mutant polymerase^18^ that can form short glycan strands but cannot crosslink them (Supplementary Fig. 4c). To visualize the polymers, we labeled the stem peptides with biotin-D-Lys (BDL) for immunoblotting^19^. No changes in the polymers were observed with LytH treatment alone, but in the presence of the complex, we saw a reduction in the intensity of BDL-labeled glycan strands (Fig. 3a and Supplementary Fig. 7b). When a radiolabel was incorporated into the glycan backbone^20^, we observed a reduction in polymer size but not in the intensity of bands (Fig. 3b and Supplementary Fig. 8). These changes, which were only observed for complexes containing catalytically-active LytH, were consistent with removal of stem peptides from the polymers. We were able to detect the tetrasaccharide-monopeptide by LC-MS analysis (Supplementary Fig. 9). We were also able to detect the stem peptide released from synthetic peptidoglycan polymers^21^ (Fig. 3c and Supplementary Fig. 10). Taken together, these experiments confirmed that the LytH complex can remove stem peptides from peptidoglycan. We have renamed the membrane protein partner ActH (<u>act</u>ivator of Lyt<u>H</u>).

**Figure 3:**
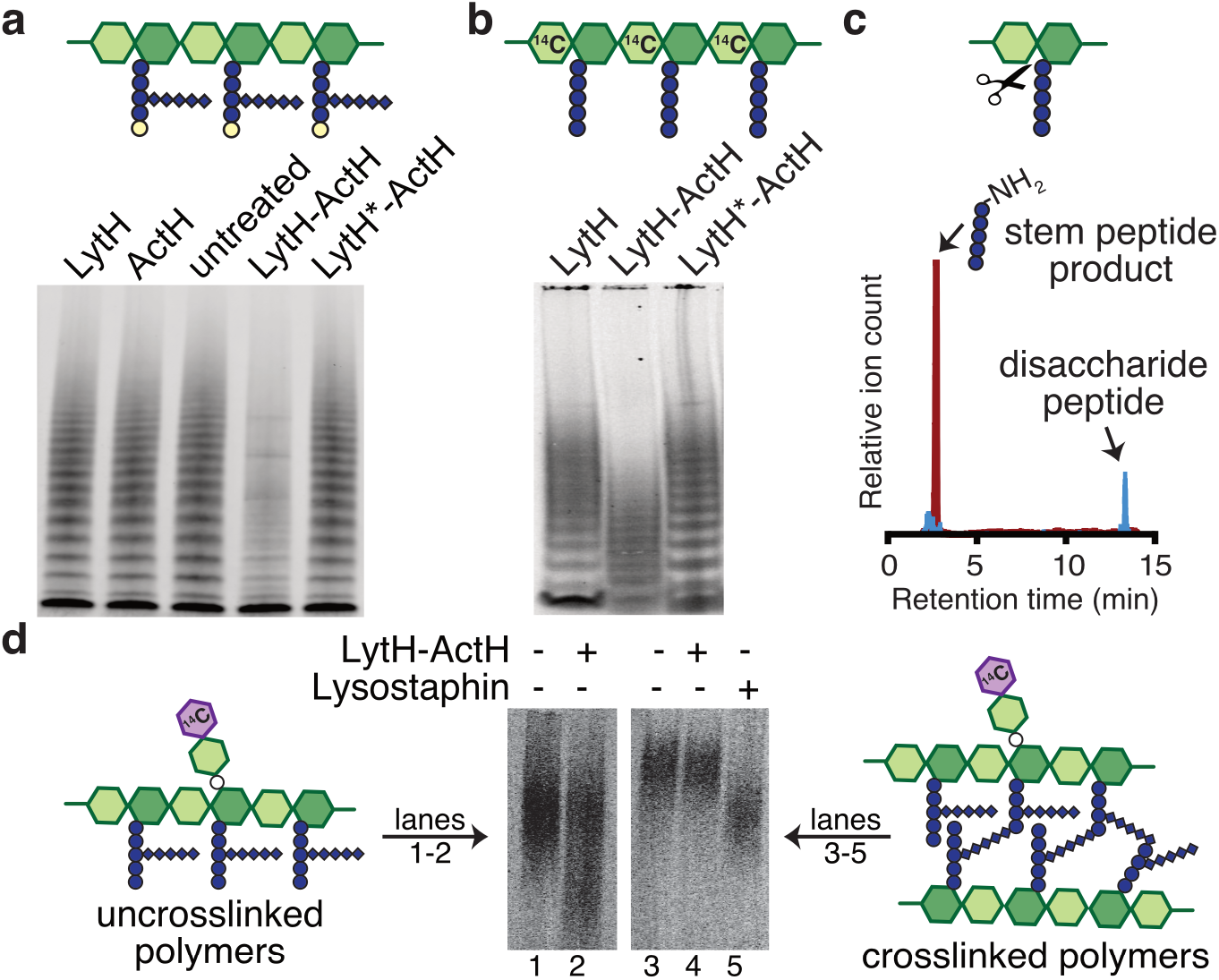
The LytH-ActH complex cleaves stem peptides from uncrosslinked peptidoglycan. **a**, LytH-ActH removes stem peptides from uncrosslinked glycan strands synthesized *in vitro*. Glycan strands were labeled with biotin-D-Lys (BDL, yellow spheres), separated by SDS-PAGE, and immunodetected with streptavidin. LytH* denotes the catalytic mutant LytH^D195A^. **b**, Uncrosslinked glycan strands containing a [^14^C]-radiolabel in the glycan backbone undergo a downward mass shift upon LytH-ActH removal of stem peptides. **c**, LC-MS analysis confirmed that LytH-ActH releases stem peptides from uncrosslinked polymers. **d**, LytH-ActH did not detectably cleave peptidoglycan crosslinks. Uncrosslinked and crosslinked polymers labeled with a [^14^C]-disaccharide on the polymer backbone were treated with the complex; a mass shift indicating cleavage was only observed for uncrosslinked polymers. The lysostaphin control cleaved crosslinks.

### LytH prefers to cleave uncrosslinked peptidoglycan

Many amidases efficiently degrade sacculi because they can act on crosslinked peptidoglycan. We tested the LytH-ActH complex with purified sacculi but did not detect hydrolytic activity, suggesting the complex prefers uncrosslinked substrates (Supplementary Fig. 11a). To directly assess substrate preferences, we needed matched substrates that only differed in whether they were crosslinked. We made uncrosslinked glycan strands from native *S. aureus* Lipid II, introduced a radiolabel on the glycan backbone, and subjected a portion of the labeled material to crosslinking^22^ (Supplementary Fig. 11b). Treatment with the LytH complex showed that the uncrosslinked material reacted to produce lower molecular weight material, but the crosslinked material did not detectably react in the presence of the complex (Fig. 3d and Supplementary Fig. 11c). We concluded that LytH is unlike previously characterized amidases involved in cell separation because it strongly prefers uncrosslinked peptidoglycan as a substrate. Consistent with the preference observed *in vitro* using synthetic peptidoglycan, we observed a substantial increase in abundance of the tetrasaccharide-monopeptide species in digests of sacculi from *lytH^WT^* cells that were treated with sublethal oxacillin to reduce crosslinking (Fig. 1e). Hence, the *in vitro* and *in vivo* evidence indicates that LytH removes stem peptides from membrane-proximal peptidoglycan that is not, or at least not extensively, crosslinked.

### Slowing peptidoglycan synthesis rescues *lytH* deletion defects

We next investigated how LytH could regulate cell division. The division defects resulting from *lytH* deletion implied LytH is important for FtsZ assembly. We found using a functional FtsZ-sGFP fusion^5^ that FtsZ is frequently mislocalized in ∆*lytH* cells (Fig. 4a and Supplementary Fig. 12). Therefore, LytH is needed for the correct positioning of FtsZ, and the divisome, at midcell. Because FtsZ assembly defects have been observed in other mutants that grow to aberrantly large size^8–10^, we speculated that the defects in divisome placement were due to uncontrolled cell growth. We exploited the observation that the ∆*lytH* mutant is temperature-sensitive to select for suppressors that might provide more information about the role of LytH (Fig. 4b). We identified multiple suppressors with distinct mutations in the polymerase domain of PBP2, a bifunctional enzyme that is the major peptidoglycan synthase in *S. aureus*^23^ (Supplementary Table 3). We found that *pbp2^WT^* overexpression was toxic in a Δ*lytH* background (Supplementary Fig. 13a). We purified one of the PBP2 suppressors, PBP2^F158L^, and found that it produced fewer and shorter glycan strands than PBP2^WT^ (Fig. 4b and Supplementary Fig. 13b,c). The *pbp2* suppressor also restored ∆*lytH* cells to WT size (Fig. 1d) and corrected FtsZ mislocalization defects (Fig. 4a). These results showed that reducing PBP2 polymerase activity compensates for loss of LytH (Supplementary Fig. 13d). If so, one would predict that moenomycin treatment, which selectively inhibits PBP2-mediated peptidoglycan polymerization^18^, would also rescue growth of the ∆*lytH* mutant at elevated temperature; indeed, this was the case (Fig. 4c). Taken together, these findings suggested that LytH controls cell growth by regulating the activity of PBP2.

**Figure 4:**
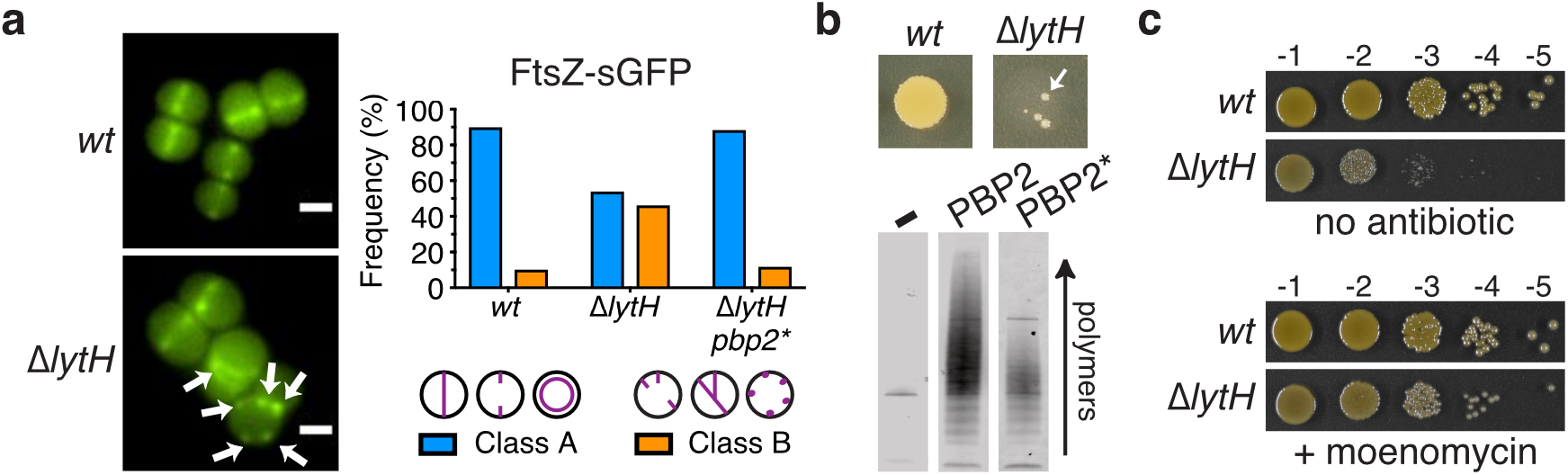
Slowing peptidoglycan synthesis compensates for loss of LytH. **a**, FtsZ was frequently mislocalized in the absence of LytH (arrows). Quantitation of FtsZ-sGFP localization (~1500 cells/strain): at midcell where a nascent or complete septum was formed (Class A); as peripheral puncta or at multiple septa (Class B). The suppressor allele *pbp2^F158L^ (pbp2*)* corrected Δ*lytH* defects. Scale bars, 1 μm. **b**, High-temperature (42°C) sensitivity of *lytH* deletion was suppressed (arrow) by reducing PBP2 activity. Purified PBP2 and a representative variant, PBP2^F158L^ (PBP2*), were incubated with Lipid II to assess polymerization activity. **c**, Reducing peptidoglycan synthesis activity pharmacologically suppressed ∆*lytH* high-temperature sensitivity. Moenomycin inhibits PBP2.

### Loss of LytH leads to excessive peripheral peptidoglycan synthesis

We used fluorescence microscopy to test whether peptidoglycan synthesis was dysregulated in the absence of LytH. To monitor sites of peptidoglycan synthesis, we first labeled cells with fluorescent D-lysine (FDL)^24^, which is exchanged into stem peptides by transpeptidase activity. Fluorescent bands at midcell reflect septal peptidoglycan synthesis. Compared with WT cells, the ∆*lytH* mutant showed fewer cells with signal at midcell and more cells with signal at misplaced septa or peripheral puncta (Fig. 5a and Supplementary Fig. 14). By measuring the ratio of fluorescence intensity at the septum vs. periphery in cells that have a single complete septum, we found that FDL-labeled stem peptides were enriched at the periphery in the absence of LytH (Fig. 5a). To determine whether the transpeptidase activity that was observed is due to reaction at sites of new peptidoglycan synthesis, we used a pulse-chase approach^25^ to selectively detect nascent peptidoglycan with fluorescent vancomycin, which binds stem peptides having terminal D-ala-D-ala residues. We found that newly synthesized peptidoglycan was enriched at the periphery in Δ*lytH* cells (Fig. 5b and Supplementary Fig. 15). Together, these experiments showed that the excess peripheral transpeptidase activity, as assessed by FDL incorporation, occurs where new peptidoglycan is being made in ∆*lytH* cells.

**Figure 5:**
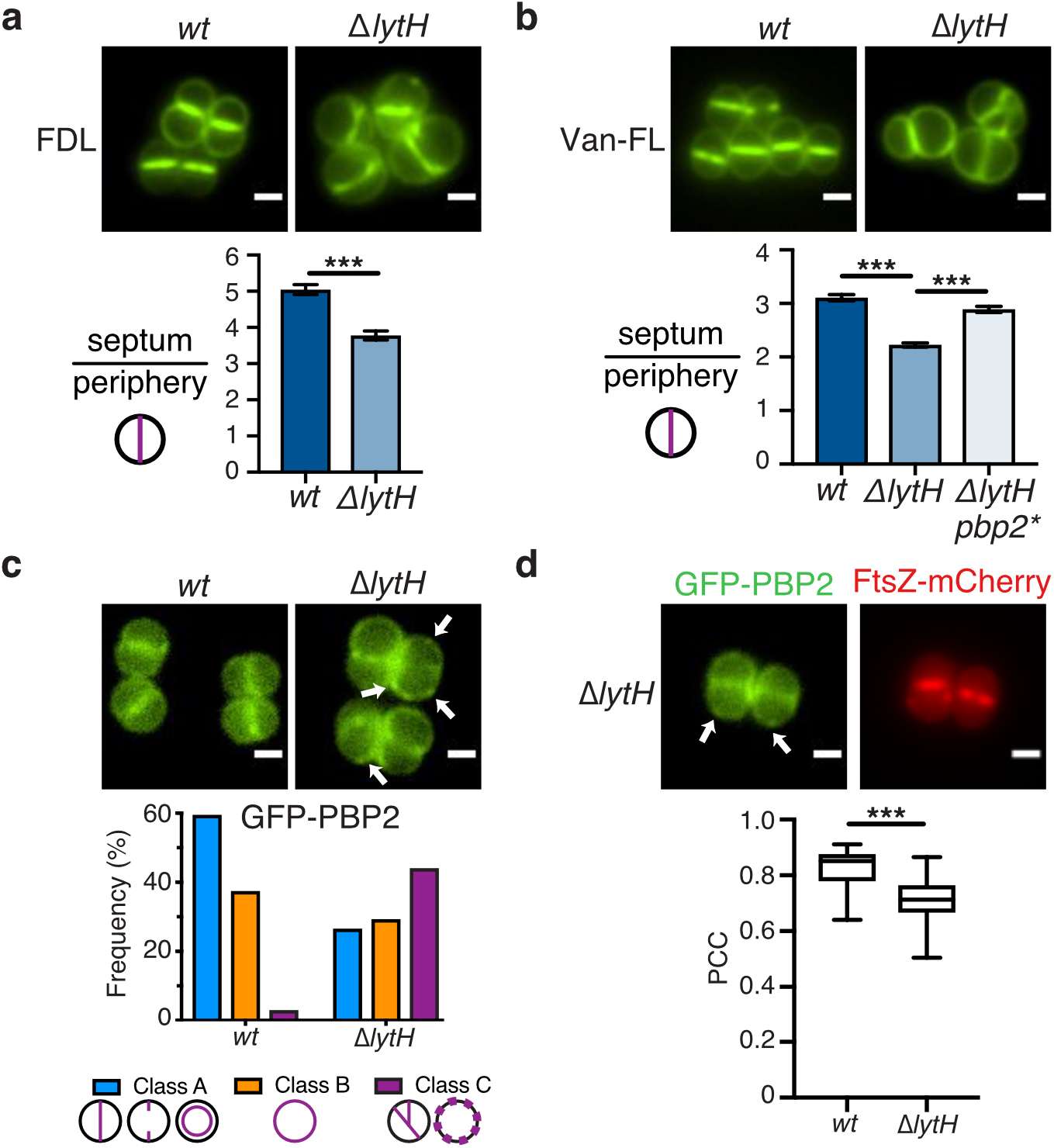
LytH regulates peripheral peptidoglycan synthesis. **a**, Cells were labeled with fluorescent D-lysine (FDL) to mark sites of transpeptidase activity; Δ*lytH* cells showed increased peripheral signal. Fluorescence ratios were calculated in cells with a single complete septum (mean ± SEM for ~160 cells/strain). All p-values were determined by two-sided Mann-Whitney U tests. \*\*\**P* < 0.001. **b**, Newly-synthesized peptidoglycan was labeled with fluorescent vancomycin (Van-FL); an increase in peripheral signal was observed for Δ*lytH* cells (mean ± SEM for ~280 cells/strain). **c**, PBP2 was frequently mislocalized in the absence of LytH (arrows). Quantitation of GFP-PBP2 localization: at midcell in dividing cells (Class A); around the membrane (Class B); at misplaced septa and/or punctate foci (Class C). Scale bars, 1 µm. **d**, GFP-PBP2 and FtsZ-mCherry showed reduced colocalization in ∆*lytH* cells: punctate PBP2 foci (arrows) were observed at sites where FtsZ was not found. Pearson correlation coefficient (PCC) data are depicted as box-and-whisker plots (~50 cells/strain). The central line indicates the median, and the bottom and top edges of each box indicate the 25^th^ and 75^th^ percentiles, respectively. \*\*\**P* < 0.001.

Our selection of *pbp2* suppressors that reduced peptidoglycan synthesis activity and corrected cell size suggested PBP2 might be responsible for the excessive peripheral peptidoglycan synthesis. Therefore, we also examined PBP2 localization using a GFP-PBP2 fusion^26^. We found that PBP2 was frequently mislocalized in ∆*lytH* cells (Fig. 5c and Supplementary Fig. 16). Mislocalization did not appear to be a consequence solely of FtsZ mislocalization because we observed some punctate PBP2 foci at peripheral regions where FtsZ was not found (Fig. 5d and Supplementary Fig. 17). Moreover, quantitative assessment showed reduced colocalization of FtsZ and PBP2. Taken together, our findings show that LytH spatially regulates peptidoglycan synthesis by reducing PBP2 activity at the cell periphery in order to control cell size and allow for proper assembly of the divisome.

## Discussion

The two key findings in this paper are the discovery of the first direct regulator for an amidase in a Gram-positive organism and the demonstration that this amidase controls cell growth. Amidases are a class of cell wall hydrolases previously shown to be important for separating daughter cells after cell division by cleaving crosslinked peptidoglycan^13^. We have shown that the amidase studied here, LytH, instead cleaves stem peptides from uncrosslinked peptidoglycan. We have also shown that LytH controls peptidoglycan synthase activity, determining how much peripheral synthesis occurs. This controls cell size and proper cell division.

A key question is how this amidase, which acts on membrane-proximal peptidoglycan, can spatially regulate peptidoglycan synthesis activity. We propose the following model (Fig. 6). Previous studies have shown that the major peptidoglycan synthase in *S. aureus*, PBP2, is recruited to midcell after FtsZ localization^5^ and recruitment of MurJ^27,28^, the flippase that exports Lipid II, the substrate for peptidoglycan synthesis, to the cell surface^5^. It has been shown that PBP2 is recruited to midcell by substrates present there with which it covalently reacts^26^. Our findings show that altering the density of PBP2 binding sites at the periphery is also necessary, as a way to control the rate of cell expansion so that the division machinery can properly localize to midcell at the appropriate time. We have found that PBP2 can react with stem peptides in preassembled glycan strands even in the absence of ongoing polymerization by the polymerase domain (Supplementary Table 4). This means that excess free stem peptides in nascent or partially crosslinked peptidoglycan could transiently trap PBP2 so that it cannot move freely to another site in the cell. Because Lipid II can diffuse freely, peripheral peptidoglycan synthesis will continue unless binding sites on peptidoglycan polymer that capture the synthase at the periphery are sufficiently reduced. Trimming of excess stem peptides reduces PBP2 activity to keep cell expansion in check. This would explain the decreased ratio of septal to peripheral synthesis even in ∆*lytH* cells that have a proper septum. According to this model, the more uncrosslinked stem peptides there are, the more important LytH is for regulating peptidoglycan synthesis to control cell size. There are substantial cell size and division defects even in cells grown at 30°C, but the morphological aberrations become more pronounced with increased temperature until LytH becomes essential for viability. LytH therefore plays a key role in controlling the spatial distribution of peptidoglycan synthases to ensure that cell expansion is coordinated with cell division.

**Figure 6:**
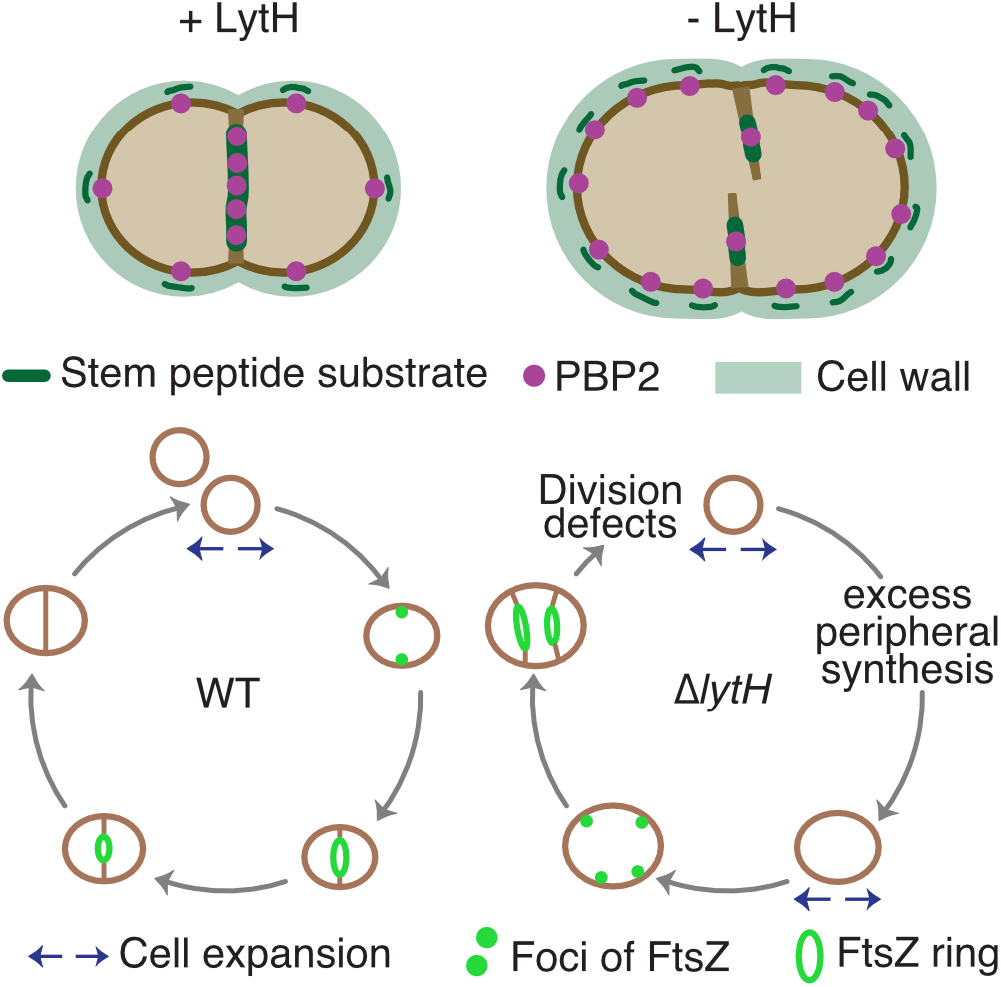
LytH coordinates cell growth with cell division. Model for LytH function. Control of cell size is required for FtsZ localization at midcell to properly initiate division. Free stem peptides limit rapid diffusion of PBP2 to midcell during division, leading to increased cell size due to excess peripheral peptidoglycan synthesis. LytH regulates stem peptide abundance on membrane-proximal peptidoglycan: in its absence, cells enlarge, FtsZ becomes mislocalized, and division initiates at aberrant sites.

We note that ActH and LytH are found in separate operons with genes important in control of protein and sugar metabolism (Supplementary Table 5), suggesting a role in adjusting peptidoglycan synthesis depending on nutrient availability. We also note that *actH* (*rbd*) forms a synthetic lethal pair with *noc*^10^, which encodes the nucleoid occlusion protein that coordinates DNA replication with division and prevents FtsZ assembly over the chromosomes. Noc has been described as a spatial regulator of cell division^7,8^. In its absence, cells grow larger than normal, resulting in division defects. The genetic connection between ActH and Noc, and the linkage between metabolism genes and *lytH*/*actH*, reinforce the importance for cell division of ensuring that cell wall synthesis is properly integrated with primary metabolic pathways inside the cell^29–31^.

The need to regulate cell wall hydrolases to control their activity in cells has long been appreciated. The peptidoglycan cell wall is essential for bacterial survival and perturbations that compromise cell wall integrity reduce cell viability. At the same time, cell wall hydrolases that cleave bonds within peptidoglycan are important for cell wall remodeling and cell separation. Hydrolytic activity must be regulated to ensure that hydrolases act when and where they should in coordination with peptidoglycan synthesis to ensure cell wall integrity^1^. Surprisingly, the first cell wall hydrolase regulators, NlpD and EnvC^32^, were only discovered recently. NlpD and EnvC are lipoproteins that activate the hydrolytic activity of cell separation amidases in Gram-negative bacteria. In the Gram-positive organisms *Streptococcus pneumoniae* and *Bacillus subtilis*, the ABC transporter-like FtsEX complex has been implicated in regulating the hydrolases PcsB^33^ and CwlO^34^, but it has not yet been established whether FtsEX directly activates these hydrolases.

Here we have shown that ActH directly interacts with LytH and activates its amidase activity. A topology prediction of ActH revealed three parts to the protein: a large intracellular region, a polytopic rhomboid fold, and an extracellular TPR domain (Supplementary Fig. 18a). Rhomboid proteins are present across all kingdoms of life, but their physiological function in bacteria is largely unknown^35^. Our work establishes a role for rhomboid proteins in bacterial cell wall biogenesis and we suggest that ActH orthologs may play a similar role in amidase activation in some other bacteria, including *B. subtilis* (Supplementary Fig. 18b). How ActH activates LytH is under investigation, although we note that ActH does not activate LytH by proteolytic processing because the amidase remains intact both in cells and *in vitro* (Supplementary Table 6). A better understanding of the domain requirements for ActH activation of LytH activity will help focus the search for additional regulators of hydrolases.

## Methods

### Bacterial growth conditions

Strains are listed in Supplementary Table 7. All *S. aureus* strains were grown in tryptic soy broth (TSB), unless otherwise indicated, containing appropriate antibiotics at temperatures ranging from 30-42°C with aeration. All *E. coli* strains were grown in Lysogeny broth (LB), unless otherwise indicated, containing appropriate antibiotics at temperatures ranging from 30-37°C with aeration. Plasmids were cloned using *E. coli* XL1-Blue, Stellar, and NovaBlue(DE3) for general purposes and *E. coli* NEB 10-beta for pTarKO constructs. Plasmids isolated from *E. coli* DC10B were directly electroporated into *S. aureus* HG003 strains without first passaging through *S. aureus* RN4220. The *E. coli* C43(DE3) strain was used for overexpression of LytH-His_6_, His_6_-ActH, His_6_-ActH-LytH^WT^-1× FLAG, and His_6_-ActH-LytH^D195A^-1× FLAG. The *E. coli* BL21(DE3) strain was used for overexpression of LytH-His_6_ truncations and PBP2-His_8_ constructs. Antibiotics were used at the following concentrations for *S.aureus* strains: kanamycin (50 μg/mL), neomycin (50 μg/mL), tetracycline (3 μg/mL), chloramphenicol (10 μg/mL), and erythromycin (10 μg/mL). Antibiotics were used at the following concentrations for *E. coli* strains: carbenicillin (100 μg/mL) and kanamycin (50 μg/mL).

### Plasmid construction

Plasmids and oligonucleotides are listed in Supplementary Tables 8 and 9, respectively. DNA sequencing of plasmids was conducted through Eton Bioscience and the Dana-Farber/Harvard Cancer Center DNA Sequencing Facility. Genomic DNA prepared from *S. aureus* strain RN4220 was used as template in PCR reactions to amplify *S. aureus* genes.

#### pTD6

The *kan^R^* marker (primers oTD3 and oTD4) and the 1-kb sequences upstream (primers oTD24 and oTD25) and downstream (primers oTD7 and oTD8) of the *lytH* open reading frame were amplified by PCR. Overlap PCR was performed to stitch together the three fragments, and the resulting fragment was cloned between the BamHI and SalI restriction sites of pKFC^36^.

#### pTP63 expression plasmids

To construct pTP63^10^ plasmids containing anhydrotetracycline (atc)-inducible constructs, the gene sequence with its native ribosome-binding site (−25) was amplified by PCR. When needed, a C-terminal 1× FLAG tag encoded in a primer was appended to the gene sequence. For plasmid pTD10, primers oTD38 and oTD39 were used. For plasmid pTD11, primers oTD40 and oTD41 were used. For plasmid pTD18, primers oTD54 and oTD58 were used. For plasmid pTD19, primers oTD56 and oTD57 were used. The gene fragments were cloned between the KpnI and BlpI restriction sites of the pTP63 vector.

#### pLOW expression plasmids

To construct pLOW^37^ plasmids containing IPTG-inducible constructs, the gene sequence with the native *rpoB* ribosome-binding site (−25) was amplified by PCR. When needed, a C-terminal 1× FLAG tag encoded in a primer was appended to the gene sequence. For plasmid pTD30, primers oTD86 and oTD85 were used. For plasmid pTD31, primers oTD87 and oTD85 were used. For plasmid pTD34, a gBlock gene fragment was synthesized by IDT. For plasmid pTD39, primers oTD86, oTD111, oTD104, and oTD106 were used; in this case, overlap PCR was performed to stitch together the two fragments. The gene fragments were cloned between the SalI and BamHI restriction sites of the pLOW vector.

#### pTD2 and pTD3

To construct plasmid pTD2 for His_6_-LytH^WT^ [E41-A291] overexpression and purification, the gene sequence for *lytH* was amplified by PCR using primers oTD15 and oTD17. To construct plasmid pTD3 for His_6_-LytH^WT^ [T102-A291] overexpression and purification, the gene sequence for *lytH* was amplified by PCR using primers oTD16 and oTD17. The gene fragments were cloned between the NdeI and BamHI restriction sites of the pET15b vector. LytH^WT^ [E41-A291] encodes a LytH truncation missing the predicted transmembrane domain. LytH^WT^ [T102-A291] encodes a LytH truncation missing the predicted transmembrane and SH3 domains.

#### pTD42

This construct is used for LytH^WT^ [M1-A291]-His_6_ overexpression and purification. The gene sequence for *lytH* was amplified by PCR using primers oTD117 and oTD116. The gene fragment was cloned between the NcoI and XhoI restriction sites of the pET28b vector.

#### pTD47

This plasmid is used to make a *tet^R^*-marked deletion of *saouhsc_01649*. The *tet^R^* marker (primers oTD145 and oTD146) and the 1-kb sequences upstream (primers oTD141 and oTD142) and downstream (primers oTD143 and oTD144) of the *saouhsc_01649* open reading frame were amplified by PCR. Overlap PCR was performed to stitch together the three fragments, and the resulting fragment was cloned between the BamHI and SalI restriction sites of the pTarKO vector^38^.

#### pTD48

This construct is for PBP2^F158L^ [K60-S716]-His_8_ overexpression and purification. The pET42a(+)-*pbp2* plasmid^17^ template was linearized by PCR using primers oTD148 and oTD149. The linearized DNA was phosphorylated at the 5’ ends using T4 polynucleotide kinase, and the ends were ligated to re-produce the circular DNA. Plasmid pET42a(+)-*pbp2^E114Q^* was constructed using QuikChange site-directed mutagenesis (StrataGene) with primers YQ1 and YQ2 and plasmid template pET42a(+)-*pbp2*.

#### pTD51

This construct is for the dual expression of His_6_-*saouhsc_01649* [N2-K487]-*lytH* [M1-A291]-1× FLAG. First, the gene sequence for *lytH* was amplified by PCR using primers oTD166 and oTD167, and the gene fragment was cloned between the NdeI and KpnI restriction sites of the pETDuet-1 vector. Next, the gene sequence for *saouhsc_01649* was amplified by PCR using primers oTD160 and oTD161, and the gene fragment was cloned between the EcoRI and HindIII restriction sites of the previously made pETDuet-1-*lytH* vector.

#### pTD52

This construct is for His_6_-*saouhsc_01649* [N2-K487] overexpression and purification. First, plasmid pTD51 was digested with NcoI and HindIII restriction enzymes. The excised NcoI-His_6_-*saouhsc_01649* [N2-K487]-HindIII fragment was then cloned between the NcoI and HindIII restriction sites of the pET28b vector.

#### pTD54

This construct is for the dual expression of His_6_-*saouhsc_01649* [N2-K487]-*lytH^D195A^* [M1-A291]-1× FLAG. pTD51 plasmid template was linearized by PCR using primers oTD103 and oTD104. The linearized DNA was phosphorylated at the 5’ ends using T4 polynucleotide kinase, and the ends were ligated to re-produce the circular DNA.

#### pTD71

This is a cadmium chloride (CdCl_2_)-inducible *S. aureus* expression vector for an FtsZ-sGFP sandwich fusion^5^. Plasmid pCN51^39^ was digested with ApaI and XhoI restriction enzymes. The excised *erm^R^* resistance cassette was then cloned between the ApaI and XhoI restriction sites of the pCN-*ftsZ^55–56^*sGFP vector.

#### pTD72

This is an anhydrotetracycline-inducible *S. aureus* expression vector for a C-terminal FtsZ [M1-R390]-mCherry fluorescent fusion with a 5-amino acid spacer sequence. This fusion protein was designed using a published sequence for a functional FtsZ-mCherry fluorescent fusion^5^. The gene sequence for *ftsZ* was amplified by PCR using primers oTD248 and oTD249. The sequence for *mCherry*^40^ was amplified by PCR from a gBlock gene fragment synthesized by IDT using primers oTD250 and oTD227. Overlap PCR was performed to stitch together the two fragments using primers oTD248 and oTD227, and the resulting fragment was cloned between the KpnI and BlpI restriction sites of the pTP63 vector.

### Strain construction

To construct the Δ*lytH*::*kan^R^* marked deletion mutant, a previously published method was used^36^. Briefly, plasmid pTD6 was electroporated into *S. aureus* strain RN4220 and transformants were selected on 10 μg/mL chloramphenicol at 30°C. Transformants were cultured at 42°C to promote chromosomal integration of the plasmid; integrants were selected on 5 μg/mL chloramphenicol at 42°C. Confirmed integrants were passaged in TSB without antibiotics at 30°C for several days to promote plasmid excision from the chromosome. Deletion mutants were selected on 50 μg/mL kanamycin and 50 μg/mL neomycin at 30°C. The Δ*lytH*::*kan^R^* deletion was confirmed by colony PCR and sequencing, and the marked deletion was transduced into other strain backgrounds to construct strains TD024 and TD240.

To construct strains containing pTP63 constructs^10^, the plasmids were first electroporated into *S. aureus* strain TD011 and transformants were selected on 10 μg/mL chloramphenicol at 30°C. The pTP63 constructs were transduced from these transformants into strain TD024 to construct strains TD036, TD037, TD074, and TD075.

To construct strains containing pLOW constructs^37^, the plasmids were first electroporated into *S. aureus* strain RN4220 and transformants were selected on 10 μg/mL erythromycin at 30°C. The pLOW constructs were transduced from these transformants into strain TD024 to construct strains TD134, TD135, TD157, and TD164.

To construct the Δ*saouhsc_01649*::*tet^R^* marked deletion mutant, a previously published method^38^ was used. Briefly, plasmid pTD47 was UV-irradiated and electroporated into *S. aureus* strain TD140, and double-crossover mutants were selected on 3 μg/mL tetracycline and 10 μg/mL targocil. The Δ*saouhsc_01649*::*tet^R^* deletion was confirmed by colony PCR, and after expressing pKFC-*tarO*, the marked deletion was transduced into strains HG003, TD024, and TD134 to construct strains TD177, TD178, and TD179, respectively.

To construct strains constitutively expressing a fluorescent GFP-PBP2 fusion, the non-replicative plasmid pPBP2-31^26^ was first electroporated into *S. aureus* strain RN4220 and transformants were selected on 10 μg/mL erythromycin at 30°C. The plasmid integrates at the native *pbp2* locus of the chromosome, and transformants were screened to identify good fluorescent colonies. The integrated pPBP2-31 construct was then transduced into strains HG003, TD024, and TD036 to construct strains TD261, TD262 and TD263, respectively.

To construct strains expressing a CdCl_2_-inducible fluorescent FtsZ-sGFP sandwich fusion^5^, plasmid pTD71 was first electroporated into *S. aureus* strain RN4220 and transformants were selected on 10 μg/mL erythromycin. From the transformants, plasmid pTD71 was transduced into strains HG003, TD024, and TD104 to construct strains TD268, TD269, and TD282, respectively. To construct dual fluorescent strains that simultaneously express GFP-PBP2^26^ and FtsZ-mCherry, plasmid pTD72 was first electroporated into *S. aureus* strain TD011 and transformants were selected on 10 μg/mL chloramphenicol at 30°C. The pTD72 construct was then transduced from the transformant into strains TD261 and TD262 to construct strains TD278 and TD279, respectively.

Phage transduction was performed using a previously published protocol^38^. Briefly, *S. aureus* donor cultures were mixed with staphylococcal phage ϕ85 and 10 mM CaCl_2_. After 30 min of incubation at room temperature, top agar (0.7% agar in TSB media) containing 7 mM CaCl_2_ was added, and the mixture was poured on tryptic soy agar (TSA) plates. The plates were incubated overnight at 30°C, and TSB was added to the plates the next day. After overnight incubation, the phage lysate was harvested from the plate and filtered. The phage lysate was used to transduce *S. aureus* recipient cultures by mixing recipient cells, phage lysate, LB media, and CaCl_2_. Transductants were selected on TSA plates containing the appropriate antibiotics and 0.05% sodium citrate. Colonies that appeared on the plates were passaged twice on sodium citrate, and transductants were confirmed using colony PCR.

### Materials

All reagents and chemicals were purchased from Sigma-Aldrich, unless otherwise indicated. Biotin-D-Lys (BDL) was prepared from Fmoc-biotin-D-lysine as previously described^41^. PCR primers were purchased from Integrated DNA Technologies. Phusion high-fidelity PCR master mix, restriction endonucleases, and T4 polynucleotide kinase were purchased from New England Biolabs, and KOD DNA polymerase was obtained from EMD Millipore. The In-fusion HD Cloning Plus kit was purchased from Takara Bio USA. Recombinant lysostaphin was purchased from AMBI Products. Mutanolysin from *Streptomyces globisporus* ATCC 21553 was purchased from Sigma-Aldrich. Genomic DNA was isolated using the Wizard Genomic DNA Purification Kit (Promega). Synthetic Lipid I was prepared as previously reported^42,43^. The radiolabeled WTA precursor, [^14^C]-LII_A_^WTA^, was prepared as previously reported^44^. Native Lipid II was extracted from *S. aureus* RN4220 culture using a published protocol^17^. Fluorescent D-lysine (FDL) was synthesized as previously reported^24^. BODIPY FL vancomycin was purchased from ThermoFisher Scientific. D-serine was purchased from Alfa Aesar.

Overexpression and purification of SgtB: *E. coli* BL21(DE3) was transformed with plasmid pET24b(+)-*sgtB^Y181D^* encoding *S. aureus* SgtB^Y181D^-His_6_ or plasmid pMgt1 encoding *S. aureus* SgtB^WT^-His_6_. These proteins were expressed and purified as reported previously^18^.

Overexpression and purification of *S. aureus* LcpB: *E. coli* BL21(DE3) was transformed with plasmid pLcpB encoding *S. aureus* His_8_-LcpB [S31-N405]. This protein was expressed and purified as previously reported^20^.

Overexpression and purification of PBPX: *E. coli* BL21(DE3) was transformed with plasmid pMW1010 encoding *E. faecalis* His_6_*-*PBPX [T36-P429]. This protein was expressed and purified as reported previously^19^.

### Transposon sequencing

A high-density transposon library of *S. aureus* HG003 was prepared^45^, treated with 37 ug/mL oxacillin, and sequenced as previously reported^11^. The oxacillin and control sequencing data can be found in the National Center for Biotechnology Information BioSample database using accession numbers SAMN08025141 and SAMN08025168, respectively. The data was processed as reported previously^11^, and genes significantly enriched and depleted under oxacillin treatment were identified using a two-sided Mann-Whitney U test corrected for multiple hypothesis testing via the Benjamini-Hochberg method. All scripts used for this analysis can be found at https://github.com/SuzanneWalkerLab/TnSeqMOAPrediction.

### Transmission electron microscopy

*S. aureus* strains were inoculated in TSB and the cultures were grown at 30°C overnight with aeration. The overnight cultures were diluted 1:100 into fresh TSB and grown to mid-log phase at temperatures ranging from 30-42°C. Cells were collected, added to an equal volume of fixative (1.25% formaldehyde, 2.5% glutaraldehyde, 0.03% picric acid in 0.1 M sodium cacodylate buffer, pH 7.4), and pelleted for the fixation process. TEM samples were prepared by the Electron Microscopy Facility at Harvard Medical School (Department of Cell Biology), and images were captured on the JEOL 1200EX instrument.

### Spot dilution assay

*S. aureus* strains were inoculated in TSB with antibiotics and the cultures were grown at 30°C overnight with aeration. Overnight cultures were diluted 1:100 into fresh TSB without antibiotics and grown to mid-log phase. The cultures were normalized, ten-fold dilutions were prepared in TSB, and 5 μL of each dilution was spotted onto TSA plates containing the appropriate antibiotics (0.1-0.125 µg/mL oxacillin or 0.01 µg/mL moenomycin A) and inducer. Plates were incubated overnight at temperatures ranging from 30-42°C. A Nikon D3400 DSLR camera fitted with an AF Micro-Nikkor 60 mm f/2.8D lens was used to take pictures of the plates.

### Muropeptide analysis

*S. aureus* strains were inoculated in TSB with antibiotics and the cultures were grown at 30°C overnight with aeration. Overnight cultures were diluted to OD_600_ = 0.02 in fresh TSB with the appropriate antibiotics and grown for 6-8 hours at 37°C. For the oxacillin treatment condition, cells were grown in TSB supplemented with 0.03125 μg/mL oxacillin. Cells were harvested (10,000 rpm for 5 min) from 2 mL of each culture. The cell pellets were processed for muropeptide analysis as previously described^46^, with the following modifications. For mutanolysin digestion, the pellet was resuspended in a 100 μL digestion buffer of 12.5 mM NaH_2_PO_4_, pH 5.5, and 10 μL of mutanolysin (4000 U/mL stock in H_2_O), and the mixture was incubated at 37°C for 16 hr on an orbital shaker (300 rpm). The lyophilized materials were resuspended in 100 μL of H_2_O and 15 μL was injected for LC-MS separation of muropeptides.

LC-MS was conducted using an Agilent Technologies 1200 series HPLC in line with an Agilent 6520 Q-TOF mass spectrometer and electrospray ionization (ESI) operating in positive ion mode and the low (1700 m/z) mass range. The Agilent Q-TOF was partially funded by the Taplin Funds for Discovery program.

The muropeptide fragments were separated on a Waters Symmetry Shield RP18 column (5 μM, 3.9 × 150 mm) with a matching column guard using previously published LC-MS conditions^17^: 0.5 mL min solvent A (H_2_O, 0.1% formic acid) for 5 min followed by a linear gradient of 0 to 40% solvent B (acetonitrile, 0.1% formic acid) over 25 min.

For targeted LC-MS/MS identification of the tetrasaccharide-monopeptide species (866.3864 [M+2H]^2+^), 25 μL of muropeptide sample from oxacillin-treated wild-type cells was injected. The same method and column as above were used, with the following modifications: prec. m/z = 866.3881; Z = 2; retention time = 14.3 min; delta retention time = 2 min; iso. width = ~4 m/z.

LC-MS data were analyzed using the Agilent MassHunter Workstation Qualitative Analysis software version B.06.00. Theoretical isotope distributions were obtained using the Thermo Xcalibur software version 3.0.63. Muropeptide species were extracted using their theoretical m/z values with a mass error of ± 20 ppm. The observed and theoretical isotope distributions were compared to confirm the species. The relative ion count for a species of interest was calculated by dividing its extracted ion count (EIC) by the sum of the EICs for the following species within the same experiment and sample (we refer to this sum as the total ion count, TIC): monomer (1253.5856 [M+H]^+^), dimer (1209.0617 [M+2H]^2+^), trimer (1194.2204 [M+3H]^3+^), and tetramer (1186.7998 [M+4H]^4+^). These are the theoretical mass-to-charge ratios used to extract the ions. All relative ion count plots represent the means and standard deviations from three independent experiments. P-values were determined using an unpaired, two-tailed *t*-test: \**P* < 0.05; \*\**P* < 0.01; ns, not significant.

### Pairwise comparison of muropeptide abundance

LC-MS total ion chromatograms were converted from Agilent Masshunter. d format to. mzdata format and uploaded to the XCMS platform (The Scripps Research Institute). Pairwise comparison of muropeptide abundance was performed with the HPLC/Q-TOF parameter. Pairwise data were sorted using the following parameters to distinguish relevant species from background signal: retention time (rtmed) between 12-18 min for this LC-MS method; fold enrichment greater than 5; maxint > 10000; and dataset1_mean > 10000.

### LytH topology prediction and homology model

LytH transmembrane topology was predicted using TMHMM Server version 2.0. LytH homology model was generated using the Phyre2 server and displayed in MacPyMol version 1.6. A sequence alignment of *S. aureus* LytH and *E. coli* AmiC was generated using Clustal Omega to identify the catalytic residues in LytH belonging to the amidase_3 family of proteins^15^.

### Overexpression and purification of full-length LytH

*E. coli* C43(DE3) was transformed with plasmid pTD42 encoding full-length *S. aureus* LytH [M1-A291] with a C-terminal His_6_ tag. Colonies were inoculated into 50 mL Terrific broth supplemented with 50 μg/mL kanamycin, and the culture was grown overnight at 37°C with aeration. The overnight culture was diluted 1:100 into 1.5 L Terrific broth supplemented with 50 μg/mL kanamycin and grown at 37°C with aeration until mid-log phase, at which point the culture was shifted to 16°C. LytH expression was induced by the addition of 1 mM isopropyl-β-D-1-thiogalactopyranoside (IPTG), and the culture was grown for another 18−20 hr at 16°C with aeration. Cells were harvested (3600 rpm, 20 min, 4°C), resuspended in 30 mL Buffer A (25 mM HEPES, pH 6.8, 200 mM NaCl, 10% glycerol) supplemented with 1 mg/mL lysozyme and 250 μg/mL DNase, and lysed using an EmulsiFlex-C3 cell disruptor (Avestin). The cell lysate was centrifuged at low speed (10000×g, 5 min, 4°C) to collect and remove cell debris, and the supernatant was transferred for ultracentrifugation (35000 rpm, 45 min, 4°C). The membrane pellet was resuspended in 30 mL Buffer A supplemented with 1% *n*-Dodecyl β-D-maltoside (DDM, Avanti Polar Lipids), homogenized with a Dounce homogenizer, and the homogenate was tumbled at 4°C for 1 hr. A second ultracentrifugation step (35000 rpm, 45 min, 4°C) was performed to remove non-homogenized membranes, and the supernatant was applied to 2 mL of pre-washed Ni-NTA resin (Qiagen) at 4°C. The Ni-NTA resin was pre-equilibrated in Buffer B (25 mM HEPES, pH 7.5, 200 mM NaCl, 10% glycerol, 1% DDM, 10 mM imidazole). After collecting the flowthrough, the resin was washed with 3 times 10 CV of Buffer A containing: 1% DDM/10 mM imidazole, 0.25% DDM/50 mM imidazole, and 0.1% DDM/80 mM imidazole. The protein was eluted with 10 CV of Buffer A supplemented with 0.05% DDM and 400 mM imidazole. The eluted LytH protein was concentrated using a 50 kD MWCO Amicon Ultra Centrifugal Filter Unit (EMD Millipore). The LytH protein was further purified using size-exclusion chromatography with a Superdex 200 Increase 10/300 GL column (GE Life Sciences) equilibrated in Buffer C (25 mM HEPES, pH 6.8, 200 mM NaCl, 10% glycerol, 0.05% DDM). The final protein concentration was measured via Nanodrop using the calculated extinction coefficient. Purified LytH protein was diluted to 20 μM in Buffer C, aliquoted, and stored at −80°C.

### Biotin-D-lysine labeling of glycan strands

A previously published method was adapted to label and detect glycan strands^19^. Briefly, 1X reaction buffer (50 mM HEPES, pH 7.5, 10 mM CaCl_2_, 60 μM Zn(OAc)_2_), BDL (3 mM), Lipid II (5 μM), and SgtB^Y181D^ (5 μM) were combined in a final volume of 10 μL (5% DMSO). The reactions were incubated at room temperature for 30 min and then quenched at 100°C for 5 min. LytH, ActH, LytH^WT^-ActH, LytH^D195A^-ActH, or LytH truncation mutants (1 μL of 20 μM stock) was added next. The reactions were incubated at room temperature for 16 hr and then quenched at 100°C for 5 min. To label glycan strands, *E. faecalis* PBPX (1 μL of 40 μM stock) was added and the reactions were incubated at room temperature for 1 hr. Finally, reactions were quenched by addition of an equal volume of 2X SDS loading buffer and analyzed using a previously published protocol^19^, with the following modifications. Five μL of each reaction was loaded into a 4−20% gradient polyacrylamide gel and run at 180V for 40 min. The products were transferred onto a 0.2 μM PVDF membrane (BioRad) and the membrane was blocked with SuperBlock TBS blocking buffer (ThermoFisher Scientific) for 1.5 hr at room temperature. Biotinylated products were probed using IRDye 800CW streptavidin (LI-COR) diluted 1:5000 in SuperBlock TBS buffer for 1 hr at room temperature. The blot was washed several times with 1X TBS, imaged using an Odyssey CLx imaging system (LI-COR), and analyzed with ImageJ.

### Co-immunoprecipitation with 1× FLAG-tagged proteins

*S. aureus* strains TD134, TD135, and TD157 carrying IPTG-inducible, FLAG-tagged full-length LytH, GFP, and truncated LytH, respectively, were grown in 25 mL TSB supplemented with the appropriate antibiotics at 30°C overnight with aeration. Overnight cultures were diluted 1:100 into 1 L TSB supplemented with 10 μg/mL erythromycin and 1 mM IPTG. Cultures were grown at 37°C for 6−8 hr with aeration to induce the expression of the FLAG-tagged proteins. Five hundred mL of cultures were harvested (3600 rpm, 15 min, 4°C) and the cell pellet was resuspended in 30 mL of lysis buffer (1X PBS, pH 7.4, 10 μg/mL DNase, 20 μg/mL RNase, 200 μg/mL lysostaphin, 5 mM MgCl_2_, 1 complete EDTA-free protease inhibitor tablet (Roche)). The resuspended cells were incubated at 37°C for 1 hr on a shaking platform to promote cell lysis, then cooled on ice, and further lysed using an EmulsiFlex-C3 cell disruptor. The lysate was spun down (10000×g, 15 min, 4°C) and the supernatant was transferred for ultracentrifugation (35000 rpm, 45 min, 4°C). The membrane pellet was resuspended in 3 mL of membrane extraction buffer (1X PBS, pH 7.4, 500 mM NaCl, 1% Triton X-100), homogenized with a Dounce homogenizer, and the homogenate was tumbled at 4°C for 1 hr. A second centrifugation step (10000×g, 15 min, 4°C) was performed to remove the remaining cell debris. The supernatant (~3 mL) was added to 3 mL of no salt wash buffer (1X PBS, pH 7.4, 1% Triton X-100) to dilute the salt. The diluted sample was batch incubated with 50 μL of pre-washed α-FLAG M2 magnetic beads (Sigma) at 4°C for 1 hr on a rocker. The α-FLAG M2 magnetic beads were pre-equilibrated in 1X PBS, pH 7.4. After incubation, the samples were transferred to a magnetic tube rack to separate the magnetic beads containing bound proteins from the supernatant. The beads were washed with 5× 500 μL of wash buffer (1X PBS, pH 7.4, 200 mM NaCl, 1% Triton X-100). To elute bound proteins, the magnetic beads were resuspended in 125 μL of elution buffer (1X PBS, pH 7.4, 200 mM NaCl, 1% Triton X-100, 150 μg/mL 3× FLAG peptide) and the samples were incubated for 15 min at room temperature to allow the proteins to elute. The tubes were then transferred to the magnetic rack and the supernatant containing eluted proteins were collected. A second elution step with 125 μL of elution buffer was performed. The eluate and wash fractions were heated at 37°C for 15 min in SDS loading buffer and then separated on a 4-20% gradient polyacrylamide gel. The gel was stained with InstantBlue Protein Stain (Expedeon) for 1 hr at room temperature, destained with H_2_O, and imaged using a FluorChem R system (ProteinSimple). Protein bands were excised and stored in deionized H_2_O prior to submission for mass spectrometry analysis at the Taplin Mass Spectrometry Facility, Harvard Medical School.

### N-terminal Edman sequencing of LytH

C-terminal FLAG-tagged LytH was expressed in strain TD134 and purified from detergent-solubilized membranes as described above. The eluate was separated on a 4-20% gradient polyacrylamide gel. Separated proteins were transferred to a 0.2 µm methanol-activated PVDF membrane. After the transfer, the membrane was washed with deionized H_2_O (5 washes, 5 min each). The membrane was then soaked in staining solution (0.02% Coomassie Brilliant Blue R-250 (ThermoFisher Scientific), 40% methanol, 5% acetic acid) for 30 sec at room temperature. The staining solution was removed and the membrane was immediately soaked in destaining solution (40% methanol, 5% acetic acid) for 1 min at room temperature. The destaining solution was removed and the membrane was washed with deionized H_2_O (6 washes, 5 min each). After the washes, the membrane was placed on a Whatman filter paper to air-dry completely. Protein bands were visible against a clear background. The LytH protein band was excised and submitted for Edman sequencing analysis at the Tufts University Core Facility.

### ActH topology prediction and phylogeny analysis

ActH transmembrane topology was predicted using TMHMM Server version 2.0. The domain structure of ActH was determined using the NCBI blastp (protein-protein BLAST) suite. Using NCBI CDART (Conserved Domain Architecture Retrieval Tool), ActH homologs with a similar domain structure as ActH, containing a rhomboid domain and tetratricopeptide repeats, were identified. Most homologs were found in the Firmicutes. Several homologs from each genus were then crosschecked using the TMHMM Server version 2.0 to confirm that their predicted membrane topologies were similar to that of ActH. The confirmed 228 CDART hits were loaded through the NCBI Batch Entrez database using their NCBI taxonomy IDs, and the 16S rRNA sequences from 196 species were extracted in FASTA format. These 16S rRNA sequences were aligned and phylogenic analysis was performed using Geneious version 9.1.5 as follows. First, a multiple sequence alignment of the 16S rRNA sequences was generated based on the ClustalW algorithm. The built-in Geneious Tree Builder function was used to generate a cladogram based on the Tamura-Nei genetic distance model and neighbor-joining method.

### Overexpression and purification of LytH-ActH complexes and ActH

*E. coli* C43(DE3) was transformed with plasmid pTD51 encoding *S. aureus* LytH^WT^-1× FLAG and His_6_-ActH, plasmid pTD54 encoding *S. aureus* Lyt^HD195A^-1× FLAG and His_6_-ActH, or plasmid pTD52 encoding *S. aureus* His_6_-ActH. Colonies were inoculated into 50 mL Terrific broth supplemented with 100 μg/mL carbenicillin, and the culture was grown overnight at 37°C with aeration. The overnight culture was diluted 1:100 into 2 × 1 L Terrific broth supplemented with 100 μg/mL carbenicillin and grown at 37°C with aeration until mid-log phase, at which point the culture was shifted to 20°C. Dual protein expression was induced by the addition of 1 mM IPTG, and the culture was grown for another 18-20 hr at 20°C with aeration. Cells were harvested (3600 rpm, 20 min, 4°C) and resuspended in 30 mL Buffer A (50 mM HEPES, pH 7.5, 500 mM NaCl, 10% glycerol) supplemented with 1 mg/mL lysozyme, 1 mM Tris(2-carboxyethyl)phosphine hydrochloride (TCEP-HCl, ThermoFisher Scientific), and 250 μg/mL DNase. The resuspended cells were stirred at 4°C for 30 min to disperse clumps and lysed using an EmulsiFlex-C3 cell disruptor. The cell lysate was centrifuged at low speed (10000 xg, 5 min, 4°C) to collect and remove cell debris, and the supernatant was transferred for ultracentrifugation (32000 rpm, 45 min, 4°C). The membrane pellet was resuspended in 30 mL of Buffer A supplemented with 1% DDM and 1 mM TCEP, and homogenized with a Dounce homogenizer. The homogenate was tumbled at 4°C for 1 hr. A second ultracentrifugation step (32000 rpm, 45 min, 4°C) was performed to remove non-homogenized membranes, and the supernatant was applied to 1 mL of pre-washed TALON metal affinity resin (Takara Clontech) at 4°C. The TALON resin was pre-equilibrated in Buffer A supplemented with 0.05% DDM and 1 mM imidazole. After collecting the flowthrough, the resin was washed with 40 CV of Buffer A containing: 1% DDM/2 mM imidazole, 0.2% DDM/4 mM imidazole, 0.1% DDM/6 mM imidazole, 0.05% DDM/8 mM imidazole, and 0.05% DDM/10 mM imidazole. The protein was eluted with 20 CV of Buffer A supplemented with 0.05% DDM and 150 mM imidazole. The eluted protein complex was concentrated using a 100 kD MWCO Amicon Ultra Centrifugal Filter Unit (EMD Millipore). The concentrated protein complex was further purified using size-exclusion chromatography with a Superose-6 Increase 10/300-GL (GE Life Sciences) column equilibrated in Buffer B (50 mM HEPES, pH 7.5, 500 mM NaCl, 10% glycerol, 0.05% DDM). The final protein concentration was measured via Nanodrop using the calculated extinction coefficient. Purified proteins were diluted to 20 μM in Buffer B, aliquoted, and stored at −80°C.

### Overexpression and purification of truncated LytH

*E. coli* BL21(DE3) was transformed with plasmid pTD2 encoding *S. aureus* His_6_-LytH^WT^ [E41-A291] or plasmid pTD3 encoding *S. aureus* His_6_-LytH^WT^ [T102-A291]. Colonies were inoculated into 25 mL LB supplemented with 100 μg/mL carbenicillin, and the culture was grown overnight at 37°C with aeration. The overnight culture was diluted 1:100 into 1.5 L LB supplemented with 100 μg/mL carbenicillin and grown at 37°C with aeration until mid-log phase, at which point the culture was shifted to 16°C. Protein expression was induced by the addition of 1 mM IPTG, and the culture was grown for another 18-20 hr at 16°C with aeration. Cells were harvested (3600 rpm, 20 min, 4°C) and resuspended in 30 mL Lysis Buffer (100 mM Tris-HCl, pH 8.0, 500 mM NaCl, 0.6% 3-((3-cholamidopropyl) dimethylammonio)-1-propanesulfonate (CHAPS, Anatrace), 0.5% Triton X-100 reduced, 100 μg/mL lysozyme, 10 μg/mL DNase, and 1 mM phenylmethylsulfonyl fluoride (PMSF)). The resuspended cells were lysed using an EmulsiFlex-C3 cell disruptor. The cell lysate was centrifuged at low speed (10000 xg, 5 min, 4°C) to collect and remove cell debris, and the supernatant was transferred for ultracentrifugation (32000 rpm, 45 min, 4°C). After ultracentrifugation, the supernatant was applied to 2 mL of pre-washed Ni-NTA resin (Qiagen) at 4°C. The Ni-NTA resin was pre-equilibrated in Buffer A (100 mM Tris-HCl, pH 8.0, 500 mM NaCl, 20% glycerol) supplemented with 2 mM imidazole. After collecting the flowthrough, the resin was washed with 20 CV of Buffer A containing: 10, 20, and 40 mM imidazole. The protein was eluted with 5 CV of Buffer A supplemented with 500 mM imidazole. The eluted protein was concentrated using a 10 kD MWCO Amicon Ultra Centrifugal Filter Unit (EMD Millipore). The concentrated protein was further purified using size-exclusion chromatography with a Superdex 75 10/300 GL column (GE Life Sciences) equilibrated in Buffer A. The final protein concentration was measured via Nanodrop using the calculated extinction coefficient. Purified proteins were diluted to 50 μM in Buffer A, aliquoted, and stored at −80°C.

### Preparation of radiolabeled peptidoglycan polymers

To prepare radiolabeled peptidoglycan oligomers, a [^14^C]-label was initially incorporated by incubating synthetic Lipid I with UDP-[^14^C]-GlcNAc (specific activity= 300 nCi/nmol) using MurG as previously published^20^. [^14^C]-Lipid II was incubated at room temperature with SgtB^Y181D^ (2 µM) in 1X TGase buffer (50 mM HEPES, pH 7.5, and 10 mM CaCl_2_) in 20% DMSO for 20 min.

To prepare WTA-modified oligomers, synthetic peptidoglycan oligomers and native peptidoglycan oligomers were first prepared cold. To prepare synthetic, cold peptidoglycan oligomers, a batch of synthetic Lipid II (20 µM) was incubated with SgtB^Y181D^ (2 µM) at room temperature in 1X TGase buffer (50 mM HEPES, pH 7.5, and 10 mM CaCl_2_) in 20% DMSO for 20 min. To prepare native peptidoglycan oligomers, a batch of native Lipid II (8 µM) was incubated with SgtB^Y181D^ (800 nM) at room temperature in 1X TGase buffer and 20% DMSO. Polymerization proceeded for 8 min before heat inactivation of SgtB^Y181D^ at 95°C for 10 min. After cooling to room temperature, the radiolabeled WTA-disaccharide precursor, [^14^C]-LII_A_^WTA^ (4.5 µM, specific activity = 300 nCi/nmol), and LcpB (2 µM) were added to each batch of peptidoglycan oligomers. Reactions proceeded for 2.5 hr at room temperature. LcpB was subsequently heat-inactivated at 95°C for 10 min.

### PAGE analysis of amidase activity

Radiolabeled peptidoglycan oligomer mixtures as prepared above were incubated with 2 µM LytH alone, LytH^D195A^-ActH, or LytH^WT^-ActH and 600 µM Zn(OAc)_2_. Reactions were incubated at room temperature overnight. Subsequently, reactions were quenched with methanol (equal volume; 10 µL) and dried to completion with a speed vacuum. Reactions were re-hydrated with 10 µL 2X SDS loading dye. The reactions were loaded on acrylamide gels, which have been described previously^47^. Tris/SDS gels (dimensions being 20 cm × 16 cm, H × W; 1.0 mM thickness) were prepared with 10 % acrylamide. The two running buffers were composed as follows: the anode buffer was 100 mM Tris (pH 8.8), and the cathode buffer consisted of 100 mM Tris (pH 8.25), 100 mM Tricine, and 0.1% SDS. The gels ran at constant voltage (30 mA) for 5 hr and were subsequently dried using a gel dryer (Labconco). Dried gels were incubated with a general phosphor screen for approximately 12-48 hr. The autoradiographs were imaged using a Typhoon 9400 imager (GE Healthcare) and analyzed using ImageJ software.

### Preparation of crosslinked WTA-oligomers

To prepare native peptidoglycan oligomers, a batch of native Lipid II (8 µM) was incubated with SgtB^Y181D^ (800 nM) at room temperature in 1X TGase buffer (50 mM HEPES, pH 7.5, and 10 mM CaCl_2_) and 20% DMSO. Polymerization proceeded for 8 min before heat inactivation of SgtB^Y181D^ at 95°C for 10 min. After cooling to room temperature, the radiolabeled WTA-disaccharide precursor, [^14^C]-LII_A_^WTA^ (4.5 µM, specific activity = 300 nCi/nmol), and LcpB (2 µM) were added to each batch of peptidoglycan oligomers. Reactions proceeded for 2.5 hr at room temperature. LcpB was subsequently heat-inactivated at 95°C for 10 min. Following precipitation and removal of both SgtB^Y181D^ and LcpB, PBP4 (2 µM) was added to each reaction and incubated at room temperature for 2 hr followed by inactivation at 95°C for 10 min. In addition to an untreated control, reactions were then subjected to either LytH^WT^-ActH (2 µM) or lysostaphin (1 mg/ml) treatment overnight at room temperature.

### Sacculi hydrolysis assay

To test if the LytH complex cleaves peptidoglycan crosslinks, hydrolysis of purified sacculi was monitored over time. Sacculi was prepared from stationary-phase cells of *S. aureus* strain HG003 using a previously published protocol^46^, with the following modifications. Volumes of solutions were scaled appropriately for the number of cells harvested. After 1 M HCl treatment, the pellets were washed with dH_2_O, flash-frozen, and lyophilized to yield purified sacculi.

A 0.5 mg/mL solution of sacculi was prepared in reaction buffer (50 mM HEPES, pH 7.5, 150 mM NaCl, 0.02% DDM, 60 μM Zn(OAc)_2_). The sacculi hydrolysis assay was set up in non-treated 96-well plates (Genesee Scientific). To each well, 150 μL of sacculi solution was added. Lysostaphin (dissolved in dH_2_O) or the purified LytH-ActH complex (stored in 50 mM HEPES, pH 7.5, 500 mM NaCl, 10% glycerol, 0.05% DDM) was mixed with the sacculi substrate. Reactions were prepared in triplicate using 0-125 nM of purified protein per reaction. The plate was covered with a lid and incubated at 25°C with shaking. Absorbance (OD_600_) was recorded at 20-min intervals for 20 hr using a SpectraMax Plus 384 microplate spectrophotometer (Molecular Devices). Hydrolysis of sacculi was monitored over time as a decrease in absorbance.

### LC-MS analysis to assess LytH activity

A previously published method was adapted to detect the products of LytH enzymatic activity by LC-MS^19^. Briefly, 1X reaction buffer (50 mM HEPES, pH 7.5, 10 mM CaCl_2_, 60 μM Zn(OAc)_2_), Lipid II (40 μM), and SgtB^WT^ (2 μM) were combined in a final volume of 10-30 μL (10% DMSO). *S. aureus* native Lipid II or synthetic Lipid II was used. The reactions were incubated at room temperature for 1 hr and then quenched at 100°C for 5 min. LytH^WT^-ActH or LytH^D195A^-ActH (2 μM) was added and the reactions were incubated at room temperature for 16 hr. The reactions were quenched at 100°C for 5 min and incubated with 4 U of mutanolysin for 1.5 hr at 37°C on an orbital shaker (300 rpm), followed by another 4 U of mutanolysin for 1.5 hr at 37°C. The muropeptides were reduced with NaBH_4_ (10-30 μL of 10 mg/mL solution in H_2_O) for 30 min at room temperature, and 20% phosphoric acid (~1.4-5 μL) was added to adjust the pH to ~4. The mixtures were lyophilized and redissolved in 20 μL of H_2_O, and 15 μL was injected for LC-MS separation using the same method as described above.

### Isolation of *lytH* suppressor mutants and whole-genome sequencing

Suppressor mutants of Δ*lytH* at 42°C on TSA spotting assay plates were independently isolated. These candidate suppressor colonies were picked and restreaked on TSA to confirm growth at 42°C. Overnight cultures of the suppressor mutants were inoculated, genomic DNA was prepared from these cultures, and the DNA concentration was measured with the Qubit dsDNA HS Assay Kit (ThermoFisher Scientific). The genomic DNA was prepared for whole-genome sequencing using a previously published method^10^, with the following modifications. Tagmented samples were pooled and analyzed by TapeStation and quantitative PCR at the Biopolymers Facility, Harvard Medical School. Sequencing was performed on the Illumina MiSeq platform using the MiSeq v3 150 cycle Reagent Kit (Illumina). Sequencing reads were analyzed using Geneious version 9.1.5. The suppressor mutants were compared to the parental *S. aureus* HG003 Δ*lytH* deletion strain to identify single nucleotide polymorphisms and deletion and insertion mutations that were unique to the suppressor mutants. The NCTC8325 genome was used as the reference genome.

### Overexpression and purification of PBP2

*E. coli* BL21(DE3) was transformed with plasmid pET42a(+)-*pbp2* encoding *S. aureus* PBP2^WT^ [K60-S716] with a C-terminal His_8_ tag or plasmid pTD48 encoding PBP2^F158L^ [K60-S716]-His_8_. These proteins were expressed and purified as reported previously^17^. PBP2^S398G^ and PBP2^E114Q^ were prepared in the same way^23^. To confirm that purified PBP2 was properly folded, a Bocillin-binding assay was used as reported previously^19^. Briefly, PBP2 (250 nM) was incubated with varying concentrations of penicillin G (1000, 100, 0 U/mL) in 1X buffer (20 mM potassium phosphate, pH 7.5, 140 mM NaCl) in a 9 μL reaction for 1 hr at 37°C on an orbital shaker (600 rpm). Bocillin-FL (1 μL of 100 μM stock) was then added and the reactions were incubated for 30 min at 37°C on an orbital shaker (600 rpm). Reactions were quenched by addition of an equal volume of 2X SDS loading buffer and analyzed by SDS-PAGE. The gel was imaged on a Typhoon FLA 9500 and analyzed using ImageJ.

### BDL-labeling assay to assess PBP2 activity

A previously published method^17^ was adapted to assess PBP2 activity. Briefly, 1X reaction buffer (50 mM HEPES, pH 7.5, 10 mM CaCl_2_), BDL (3 mM), Lipid II (10 μM), and PBP2 (1 μM) were combined in a final volume of 10 μL (10% DMSO). The reaction was incubated at room temperature for 5-15 min and then heat-quenched at 100°C for 1 min. As appropriate, lysostaphin was added to resolve highly-crosslinked peptidoglycan: 0.5 μL of lysostaphin (1 mg/mL stock) was added and the reactions were incubated at 37°C for 3 hr on an orbital shaker (300 rpm). Reactions were quenched by addition of an equal volume of 2X SDS loading buffer, and analyzed by SDS-PAGE and western blotting analysis with streptavidin as described above.

### LC-MS analysis of crosslinked muropeptides by different polymerases

Native *S. aureus* Lipid II (25 µM) was incubated with i. SgtB (1 µM), ii. the transpeptidase-inactive PBP2^S398G^ (1 µM), or iii. PBP2^WT^ (2 µM) in a 20 µl reaction containing 20% DMSO and 1X reaction buffer (50 mM HEPES, pH 6.5, 2.5 mM MgCl_2_). For i. and ii., the polymerase was heat-inactivated at 95°C for 5 min after a 2-hr incubation at room temperature; after cooling to room temperature, the glycosyltransferase-inactive PBP2^E114Q^ (1 µM) was added and the reactions were incubated at room temperature for an additional 2 hr alongside reaction iii. All reactions were then heat-inactivated at 95°C for 5 min. After cooling to room temperature, mutanolysin (1 U) was added to each reaction for 1.5 hr at 37°C followed by another 1 U aliquot for 1.5 hr. The cleaved disaccharide fragments were then reduced with NaBH_4_ treatment (10 µL of 10 mg/mL solution in H_2_O) for 30 minutes. Phosphoric acid (20%, 1.4 µL) was added to each reaction to adjust the pH to 4, and then the mixture was lyophilized and re-dissolved in 20 µl H_2_O for LC/MS analysis, conducted with ESI-MS operating in positive mode on a Bruker qTOF mass spectrometer. The same column and method as for muropeptide analysis were used.

### Fluorescent vancomycin labeling to measure cell size

A previously published method^10^ was adapted for fluorescent vancomycin (Van-FL) labeling of live *S. aureus* cells for fluorescence microscopy. Overnight cultures were diluted to a starting OD_600_ of 0.02 in 20 mL TSB, grown at 37°C with aeration to mid-log phase, and normalized to the same OD_600_. To 1 mL of cells, 1 µL of a 1:1 (v/v) mixture of 0.5 mg/mL fluorescent vancomycin and 0.5 mg/mL vancomycin-hydrochloride was added. The cells were labeled for 5 min at room temperature, with foil covering the tubes to protect the reactions from light. The cells were centrifuged (4000 xg, 1 min, 25°C), washed once with 200 µL 1X PBS, pH 7.4, and resuspended in 75-100 µL of 1X PBS, pH 7.4. Cell volumes were calculated using only non-dividing cells without a visible septum.

### Fluorescent D-lysine labeling to visualize sites of transpeptidase activity

A previously published method^41^ was adapted for fluorescent D-lysine (FDL) labeling of live *S. aureus* cells for fluorescence microscopy. Overnight cultures were diluted to a starting OD_600_ of 0.02 in 20 mL TSB, grown at 37°C with aeration to mid-log phase, and normalized to the same OD_600_. One mL of culture was centrifuged (4000 xg, 1 min, 25°C), and the cell pellet was resuspended in 500 μL of TSB. To the resuspended cells, 100 μM FDL was added and the cells were labeled for 10 min at 37°C on an orbital shaker (300 rpm), with foil covering the tubes to protect the reactions from light. The cells were centrifuged (4000 xg, 1 min, 25°C), washed twice with 200 μL 1X PBS, pH 7.4, and resuspended in 50 μL of 1X PBS, pH 7.4.

### Labeling of nascent peptidoglycan

A published method^25^ was used to label newly synthesized peptidoglycan in live *S. aureus* cells for fluorescence microscopy. Strains were grown overnight in TSB containing excess D-serine (0.125 M). Overnight cultures were diluted to a starting OD_600_ of 0.02 in 3 mL of the same medium and grown at 37°C with aeration to mid-log phase. All 3 mL of cells were centrifuged (5000 xg, 1 min, 25°C), washed once with 200 μL of TSB, and resuspended in 3 mL of TSB without D-serine supplemented. Cultures were then incubated for 20 min on a shaker at room temperature to allow incorporation of D-alanine into the cell wall. One to two mL of culture were centrifuged (4000 xg, 1 min, 25°C), and the cell pellet was resuspended in 500 μL of TSB. To the resuspended cells, 0.5 µL of a 1:1 (v/v) mixture of 0.15 mg/mL fluorescent vancomycin and 0.15 mg/mL vancomycin-hydrochloride was added. The cells were labeled for 5 min at room temperature, with foil covering the tubes to protect from light. The cells were centrifuged (4000 xg, 1 min, 25°C), washed once with 200 µL 1X PBS, pH 7.4, and resuspended in 50 µL of 1X PBS, pH 7.4.

### Visualization of fluorescent fusion proteins

To assess GFP-PBP2 localization, the strains TD261, TD262, and TD263 were used. To assess FtsZ^55–56^sGFP localization, the strains TD268, TD269, and TD282 were used. To assess GFP-PBP2 and FtsZ-mCherry colocalization, the strains TD278 and TD279 were used. All strains were grown in TSB overnight; 10 μg/mL erythromycin was added for FtsZ^55–56^sGFP strains to maintain the plasmid. Overnight cultures were diluted to a starting OD_600_ of 0.02 in 3 mL TSB. For strains TD263, TD278, and TD279, the medium was supplemented with 0.4 μM of the inducer anhydrotetracycline. For FtsZ^55–56^sGFP strains^5^, the medium was supplemented with 10 μg/mL erythromycin and 0.1 μM of the inducer CdCl_2_. Cultures were grown at 37°C or 42°C with aeration to mid-log phase. One to two mL of cells were centrifuged (4000 xg, 1 min, 25°C), washed once with 200 μL 1X PBS, pH 7.4, and resuspended in 50 μL of 1X PBS, pH 7.4.

### Wide-field epifluorescence microscopy

Cells were spotted onto a thin 2% agarose pad prepared using 1X PBS, pH 7.4. A 1.5 cover slip was used and Valap sealant (equal weight of petroleum jelly, lanolin, and paraffin) was applied to seal all sides of the cover slip. Cells were imaged using brightfield, phase-contrast, and wide-field epifluorescence microscopy at the MicRoN (Microscopy Resources On the North Quad) facility, Harvard Medical School. Images were obtained using a Nikon Ti inverted microscope fitted with a custom-made cage incubator set at 30°C, a Nikon motorized stage with an OkoLab gas incubator and a slide insert attachment, an Andor Zyla 4.2 Plus sCMOS camera, Lumencore SpectraX LED Illumination, Plan Apo lambda 100×/1.45 Oil Ph3 DM objective lens, and Nikon Elements 4.30 acquisition software. The microscope was fitted with Chroma ET filter cubes in a motorized filter turret: DAPI (49000), CFP (49001), GFP (49002), YFP (49003), and mCherry (49008). The following exposure times were used: 50 ms (FDL labeling), 100-200 ms (Van-FL labeling), 100 ms (FtsZ-mCherry), and 5 s (GFP-PBP2 and FtsZ^55–56^sGFP). Images were analyzed using the software FIJI and MATLAB scripts developed in-house.

### Semi-automated image analysis

*S. aureus* cell volumes were estimated using StaphSizer, a home-built MATLAB code, which was also used to categorize cellular phenotypes. Segmentation of cell clusters was performed on brightfield images using a watershedding algorithm^48^. First, brightfield images were used to create a binary image, to which a Euclidean distance transform function^49–51^ was applied to obtain an image whose centers corresponded to the cell regions within clusters of cells. Local minima that are not centered within the cells in the distance transform image were filtered out to prevent oversegmentation. Finally, the watershed algorithm was applied to the modified distance transform image to obtain the ridge lines that define the borders between neighboring cells. From this segmented image output, the x-y coordinates of each pixel on the cell boundary were obtained.

To sort cells based on phenotypes, the calculated cell boundary was overlaid on top of the Van-FL fluorescence image and paired with its corresponding brightfield image to enable visual inspection of each cell. A 2-dimensional ellipse function was fitted to each cell boundary to calculate the lengths of the major and minor axes of the ellipse. To estimate the volume, *S. aureus* geometry was modeled as a prolate spheroid^52^ with the following volume equation:

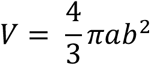

Where V is the volume, a is half of the major (longer) axis, and b is half of the minor (shorter) axis of each fitted cell. Cell volumes were calculated using approximately 700 non-dividing cells without a septum. P-values were determined by two-sided Mann-Whitney U tests.

The ratio of fluorescence intensity at the septum vs. periphery (F_septum_/F_periphery_) for selected cells was calculated using FreehandPeriSep, a home-built MATLAB code. After background correction, the coordinates of the boundary of a cell with a single complete septum were defined by the user manually tracing on a phase-contrast image. The coordinates of the user-defined region of interest (ROI) were used to create a binary mask that encompasses the entire cell (outer ROI). To obtain the fluorescence signal at the septum (septum ROI) in the same cell, the user manually traces the septum on the corresponding fluorescence image. To obtain the fluorescence signal at the cell periphery (periphery ROI), an inner ROI containing the septum and cytoplasm in the cell was first obtained by eroding the original binary mask such that the peripheral region is excluded; this inner ROI was then subtracted from the outer ROI. Finally, the periphery and septum ROIs were used to retrieve all pixel values at the cell periphery and septum, respectively. Only 50% of the brightest pixels at the septum and at the cell periphery were considered to avoid inclusion of misidentified pixels. Fluorescence ratios were calculated with the following equation, and p-values were determined by two-sided Mann-Whitney U tests^5^:

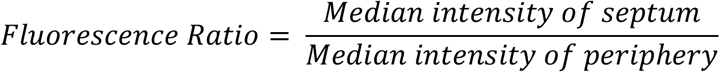

The Pearson correlation coefficient (PCC)^5^ between two fluorescence channels was calculated using FreehandPCC, a home-built MATLAB code. A higher PCC indicates a greater degree of colocalization between GFP-PBP2 and FtsZ-mCherry in a strain. Only cells showing FtsZ signal at the septum were considered for PCC analysis. The coordinates of the boundary of a cell were defined by the user manually tracing on a phase-contrast image. These coordinates were used to isolate images of chosen cells from the two background-corrected fluorescence channels. The PCC between the channels were obtained using the built-in MATLAB function *corr2*, which calculates the 2-D correlation coefficient of two images of size *m*-by-*n* with the following equation:

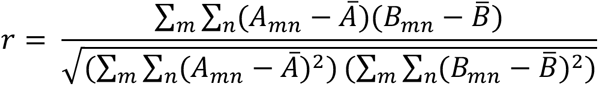

Where *A*_*mn*_ and *B*_*mn*_ are the pixel intensities for the two fluorescence channels, and 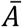 and 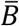 are the mean intensities of those channels. Data are represented as box-and-whisker plots in which the line inside the box indicates the median PCC, and the bottom and top edges of the box indicate the 25^th^ and 75^th^ percentiles, respectively. Whiskers are drawn to the minimum and maximum PCC. P-values were determined by two-sided Mann-Whitney U tests.

## Data availability

Transposon sequencing data (accession nos. SAMN08025141 and SAMN08025168) can be found in the NCBI BioSample database. Whole-genome sequencing data will be posted to the NCBI BioSample database. All other data are available in the manuscript or the supplementary materials. Code will be available at https://github.com/SuzanneWalkerLab/imageanalysis.

## Acknowledgements

We thank M. Welsh and A. Taguchi for help with LC-MS analysis and protein expression. We thank T. Pang from the Bernhardt and Rudner labs for strain TD215. Fluorescence images were partially acquired at the Microscopy Resources on the North Quad (MicRoN) core at Harvard Medical School. We acknowledge support from the NSF (DGE1144152 to T.D.), the NIH (P01AI083214, R01GM076710, R01AI139011, R01AI099144), and the European Research Council (ERC-2017-CoG-771709).

## Author contributions

S.W. and T.D. designed experiments and analyzed the data with advice from M.G.P on acquisition and analysis of fluorescence images. T.D. performed all experiments except the radiolabeled gel assays, which were performed by K.S, and some microscopy experiments, which were performed by P.B.F. A.G.S. and T.D. developed scripts for image analysis. K.C. analyzed transposon sequencing data. S.W., D.K., and T.D. wrote the manuscript with input from all authors.

