## Supplementary Information for "The cell cycle in *Staphylococcus aureus* is regulated by an amidase that controls peptidoglycan synthesis"

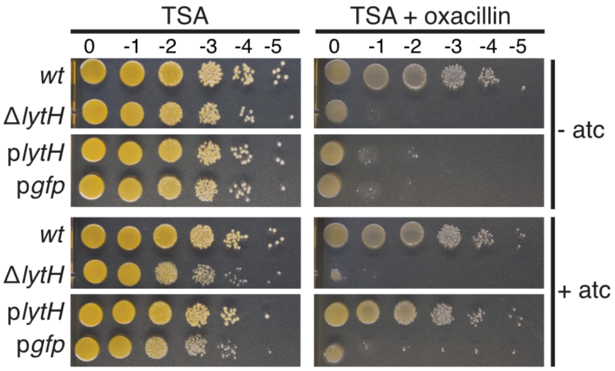


**Supplementary Figure 1:** Deletion of *lytH* sensitizes cells to oxacillin and causes septal defects. HG003 strains were spotted on TSA containing 0.1 µg/mL oxacillin and 0.4 µM of the inducer, anhydrotetracycline (atc), when indicated. Strains expressing *lytH* or *gfp* from an atc-inducible promoter in a *lytH* deletion background are denoted by p*lytH* or *pgfp*, respectively. Plates were incubated overnight at 37°C. The remaining strains that were tested in this same experiment are shown in Supplementary Fig. 6.

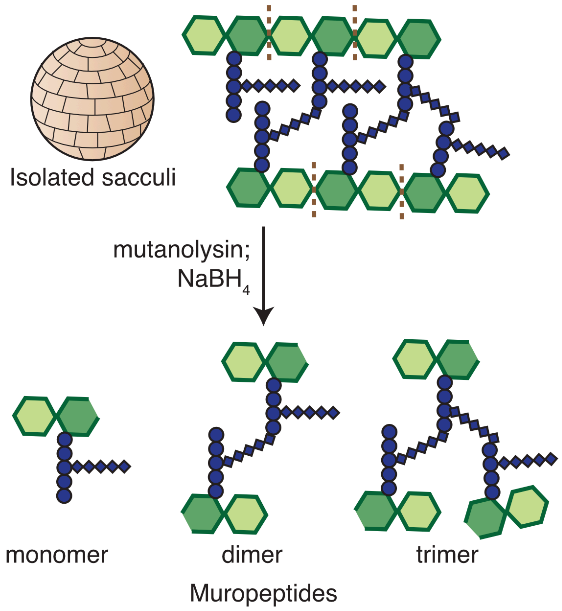


**Supplementary Figure 2:** Schematic of sacculi isolation and mutanolysin/$NaBH_4$ treatment to generate muropeptides for LC-MS analysis. The legend is the same as in Fig. 1a.

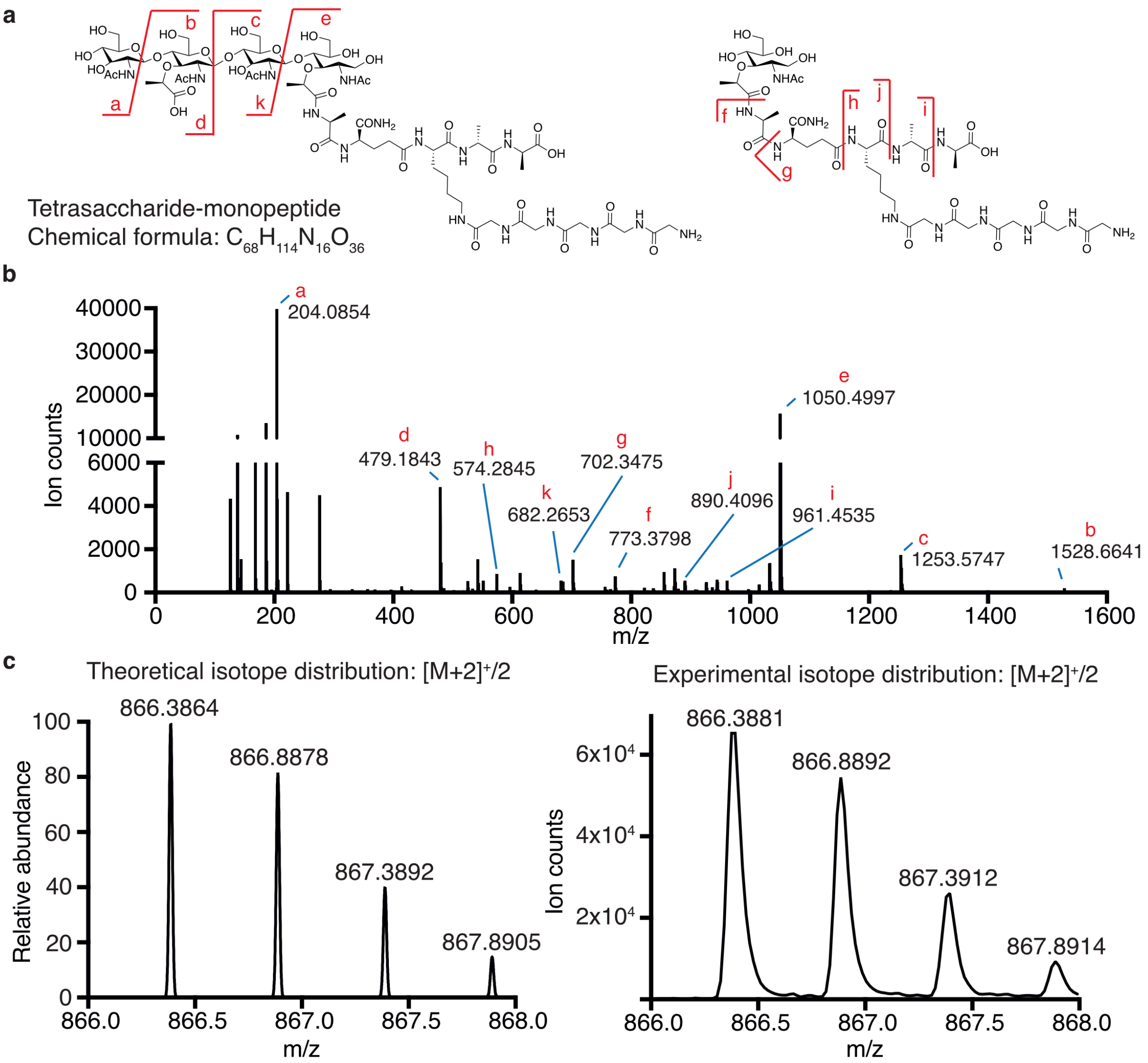


**Supplementary Figure 3:** Targeted MS/MS of the tetrasaccharide-monopeptide species. (a) Species X was enriched in sacculi isolated from *S. aureus* cells expressing *lytH*$^{WT}$. A chemical structure consistent with this species, based on an $[M]^{+2}$ ion with experimental m/z = 866.3881, is shown on the left. (b) The $[M]^{+2}$ ion was targeted for fragmentation to confirm species X was a tetrasaccharide-monopeptide. (c) The theoretical and experimental isotope distributions for the tetrasaccharide-monopeptide species are shown.

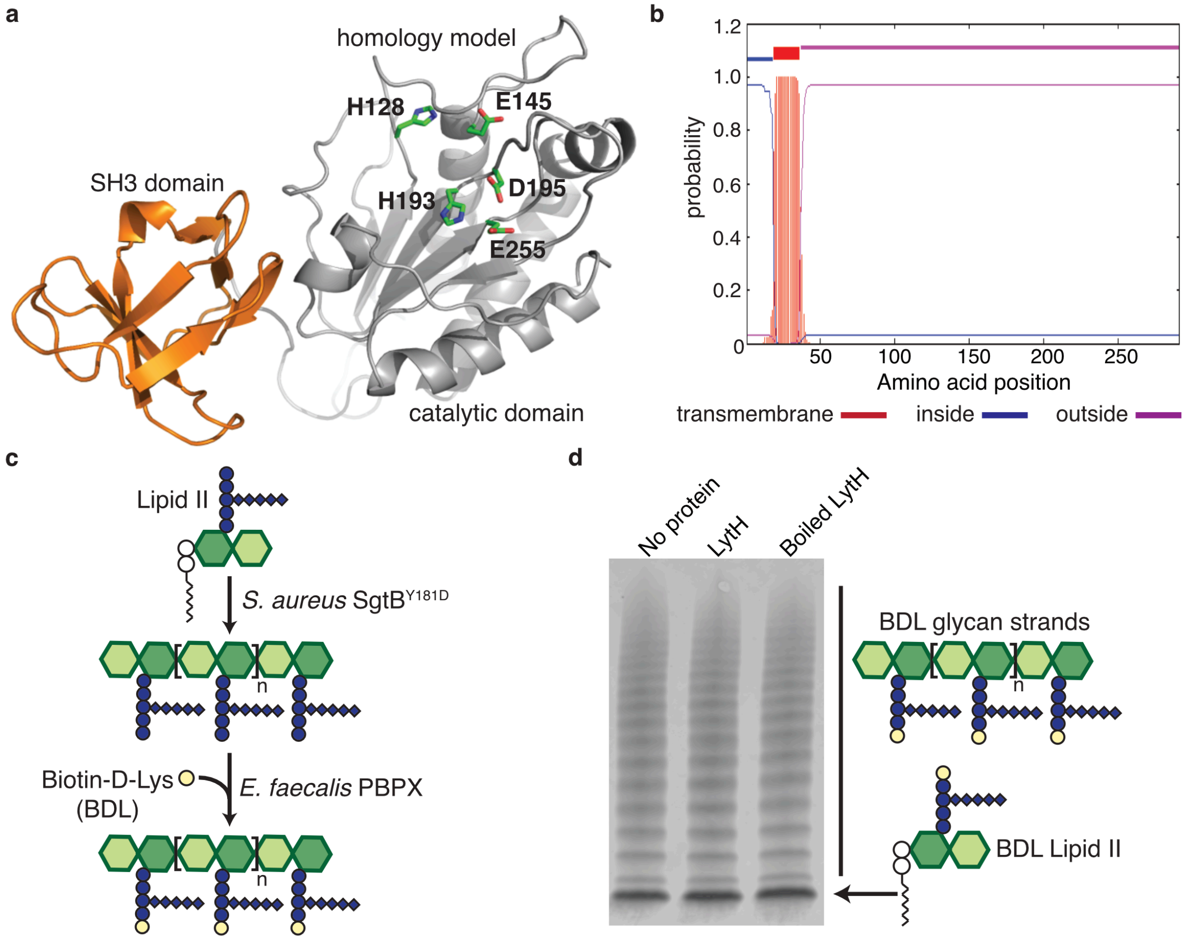


**Supplementary Figure 4:** LytH alone is not active *in vitro*. (a) A homology model of LytH highlights the predicted catalytic residues identified by sequence alignment of *S. aureus* LytH with *E. coli* AmiC[1]. All catalytic residues required for amidase activity are intact. (b) TMHMM topology prediction suggests that LytH is anchored in the cytoplasmic membrane. (c) Schematic representation of the BDL-labeling assay to assess LytH hydrolytic activity *in vitro*. Short glycan strands were synthesized from Lipid II using the processivity-defective SgtB$^{Y181D}$ glycosyltransferase[2]. *E. faecalis* PBPX[3] was used to exchange the terminal amino acid of stem peptides for biotin-D-Lys (BDL, yellow spheres), enabling visualization of glycan strands by western blotting with streptavidin. (d) Glycan strands synthesized *in vitro* were treated with LytH, resolved by SDS-PAGE, and detected by western blotting with streptavidin. No difference in the distribution or intensity of polymers was observed between the three lanes.

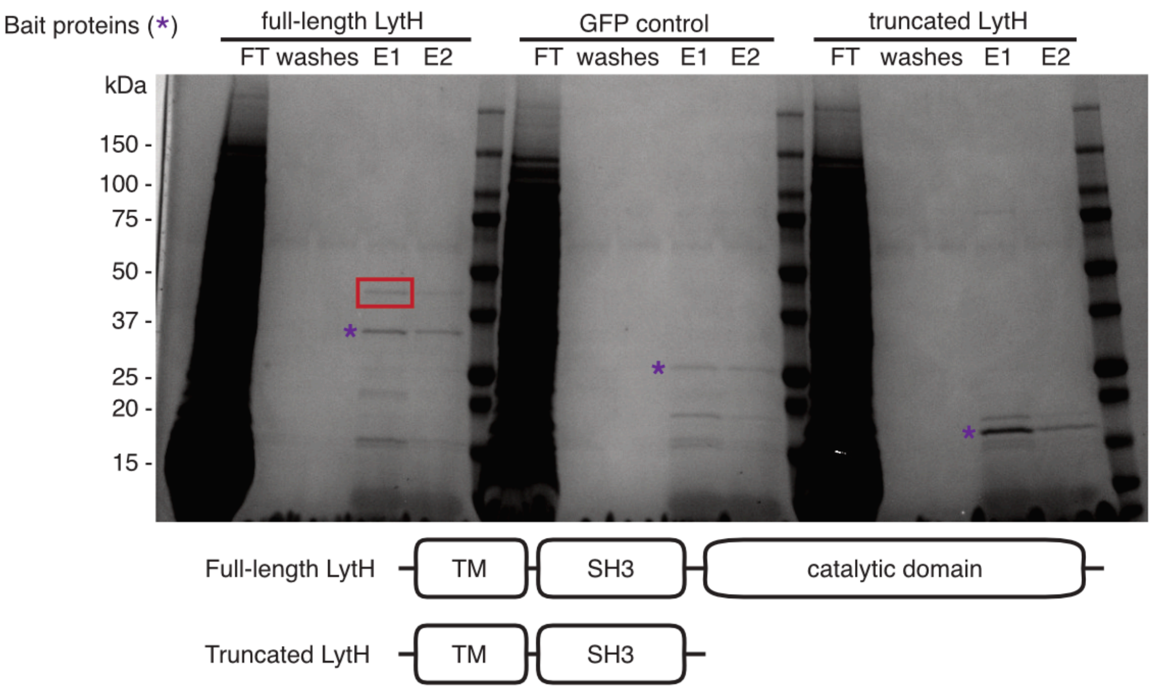


**Supplementary Figure 5:** LytH pulled down ActH, a previously uncharacterized polytopic membrane protein. SDS-PAGE analysis of the co-immunoprecipitation experiment to identify proteins that bound to full-length LytH. The C-terminal 1x FLAG-tagged proteins were expressed in *S. aureus* strain HG003 Δ*lytH*, and FLAG-tagged proteins and their partners were isolated from detergent-solubilized membranes. Purple asterisks mark the protein bands corresponding to the bait proteins. A red box is drawn around the ActH protein band. ActH is encoded by the gene *saouhsc_01649*. The fractions are: (FT) flowthrough and (E1-E2) elutions 1-2. The schematics for full-length and C-terminal truncated LytH constructs are shown with the predicted transmembrane (TM), SH3, and amidase catalytic domains annotated. This is the full gel for Fig. 2a.

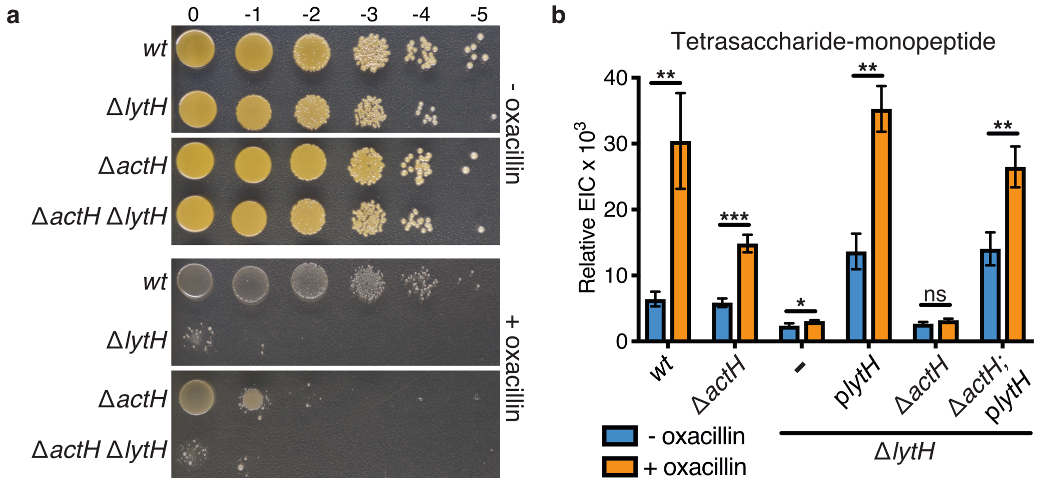


**Supplementary Figure 6:** Deletion of *actH* phenocopies deletion of *lytH*. (a) HG003 strains were spotted on TSA ± 0.125 µg/mL oxacillin, and plates were incubated overnight at 37°C. The remaining strains that were tested in this same experiment are shown in Supplementary Fig. 1. (b) The relative ion count (extracted ion count/total ion count) for the tetrasaccharide-monopeptide was calculated as described in Materials and Methods. For ease of visualization, relative ion counts were multiplied by $10^3$. Data represent the mean ± standard deviation from three independent experiments. P-values were determined using unpaired, two-tailed *t*-tests: **P* < 0.05; ***P* < 0.01; ns, not significant.

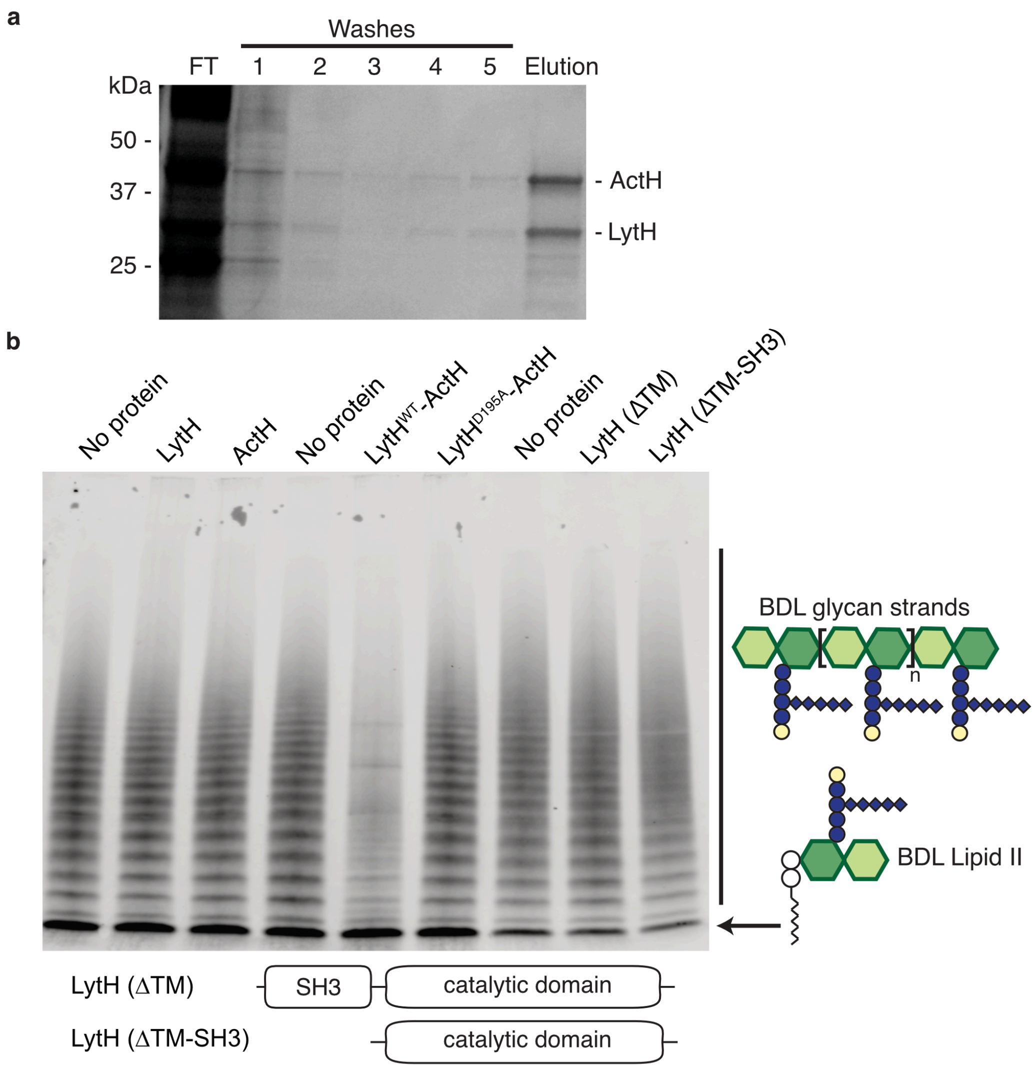


**Supplementary Figure 7:** The LytH-ActH complex removes stem peptides from glycan strands. (a) Full-length LytH and ActH were co-expressed in *E. coli* and purified from detergent-solubilized membranes as a stable complex. ActH contains an N-terminal hexahistidine tag. SDS-PAGE analysis of fractions from TALON purification of the $LytH^{WT}$-ActH protein complex: (FT) flowthrough. (b) Western blot analysis of LytH reactions with glycan strand substrate, showing a distinct difference for the unboiled, folded WT complex. Glycan strands were made *in vitro* from native *S. aureus* Lipid II and treated with the indicated proteins: LytH, ActH, $LytH^{WT}$-ActH, $LytH^{D195A}$-ActH, and LytH truncation mutants. To visualize the glycan strands and assess reaction, the strands were labeled with BDL using PBPX. For a schematic of the reaction setup, refer to Supplementary Fig. 4c. This is the full blot for Fig. 3A.

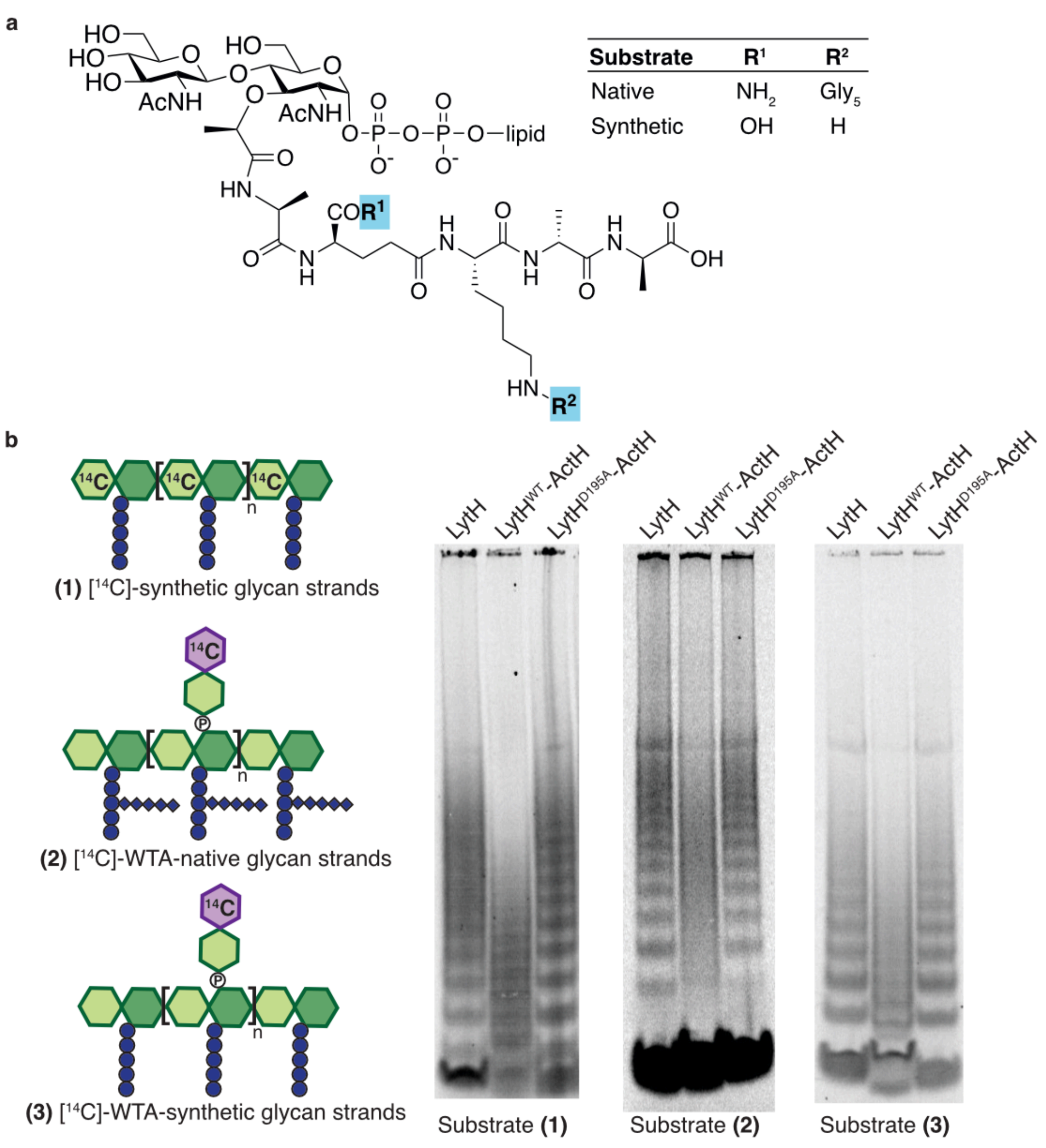


**Supplementary Figure 8:** The LytH-ActH complex removes stem peptides from radiolabeled glycan strands. (a) Chemical structure highlighting the differences between native and synthetic Lipid II. These precursors were polymerized by SgtB to make uncrosslinked glycopolymers. (b) PAGE autoradiographs of reactions with different uncrosslinked glycan strand substrates (left) in the presence of LytH, $LytH^{WT}$-ActH, or $LytH^{D195A}$-ActH. The glycopolymers for substrates (1) and (3) were made from synthetic Lipid II, while the glycopolymers for substrate 2 were generated from native Lipid II. For substrate (1), the radiolabel was found in the N-acetylglucosamine residues ([$^{14}$C]-GlcNAc) of the glycan backbone; this gel is also shown in Fig. 3b. For substrates (2) and (3), the radiolabel was provided by a short wall teichoic acid (WTA) moiety ([$^{14}$C]-$LII_A^{WTA}$) attached to the hydroxyl group at the C6 position[4,5].

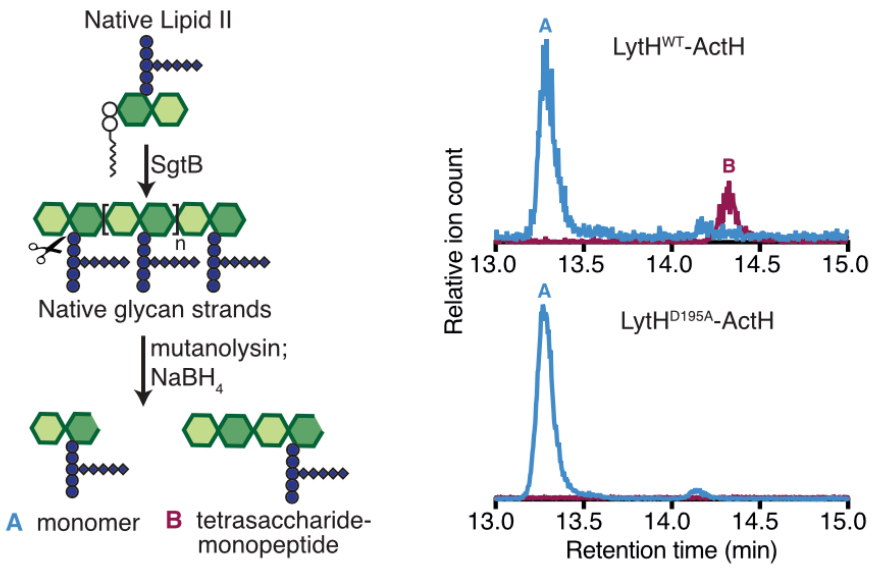


**Supplementary Figure 9:** The LytH-ActH complex releases stem peptides from glycan strands. (Left) Glycan strands were made from native Lipid II using SgtB$^{WT}$ protein and treated with the LytH complex prior to LC-MS analysis as above. (Right) LC-MS traces for the LytH *in vitro* reactions using native glycan strands. In the presence of LytH$^{WT}$-ActH, the tetrasaccharide-monopeptide was observed, but only the monomer (starting material) was observed in the LytH$^{D195A}$-ActH reaction. We could not detect the released stem peptide in these reactions with native glycan strands likely due to the poor ionization efficiency of the released pentaglycine-containing stem peptide.

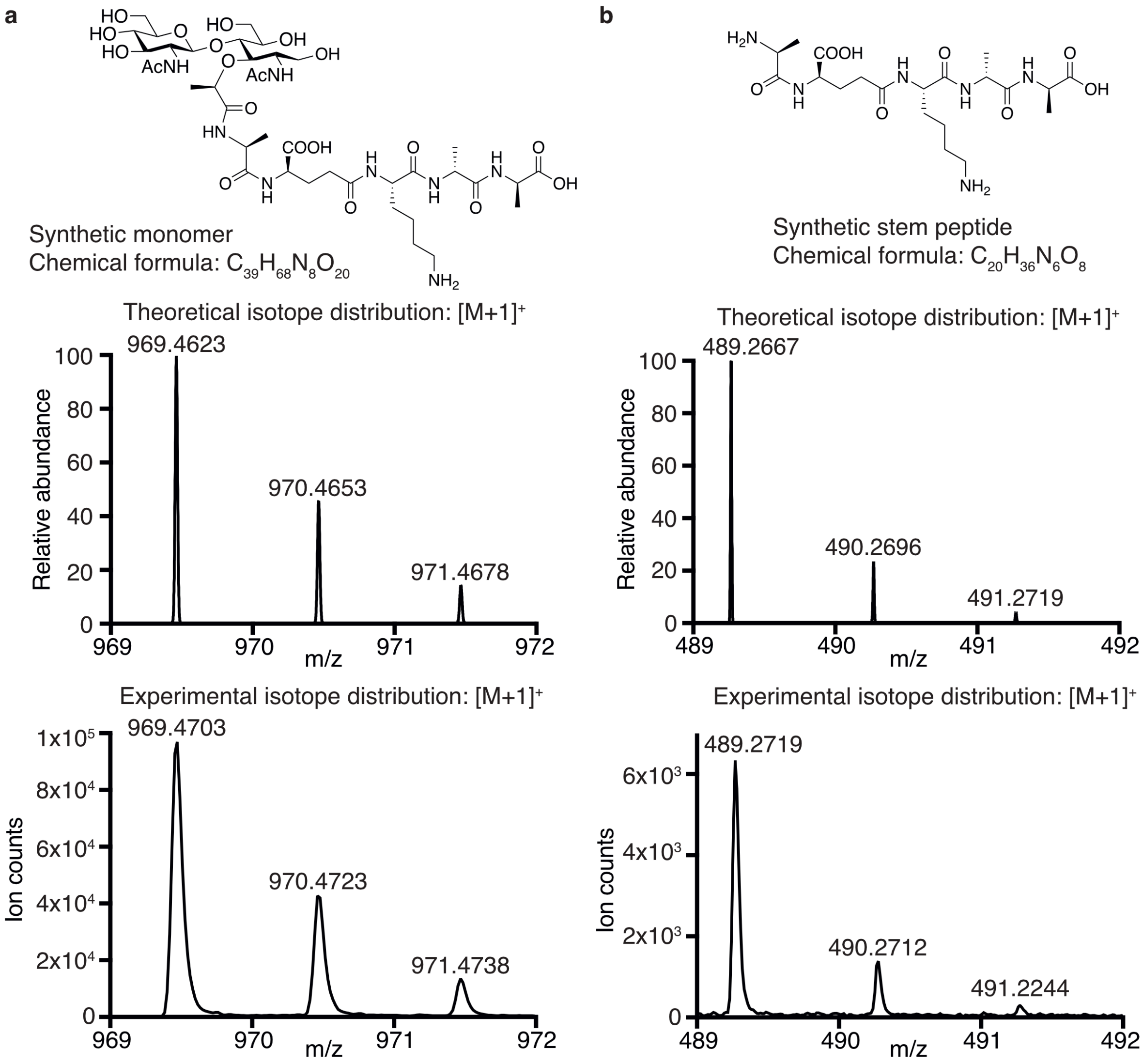


**Supplementary Figure 10:** Isotope distributions for the LytH-ActH reaction products. (a) The chemical structure and theoretical/experimental isotope distributions for the disaccharide peptide species (blue peak) shown in Fig. 3c. (b) The chemical structure and theoretical/experimental isotope distributions for the synthetic stem peptide product (red peak) shown in Fig. 3c.

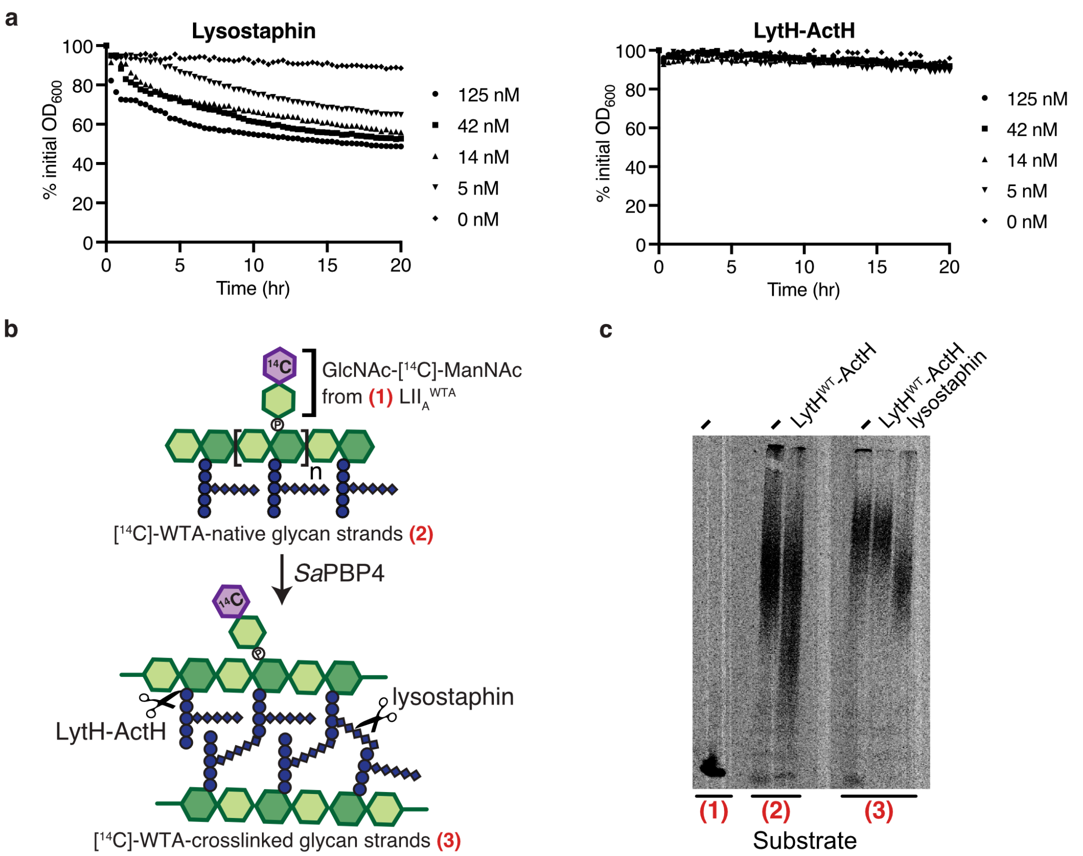


**Supplementary Figure 11:** LytH-ActH does not detectably cleave peptidoglycan crosslinks. (a) Purified *S. aureus* sacculi were treated with different concentrations of lysostaphin and LytH$^{WT}$-ActH. Hydrolysis of sacculi was monitored over time as a decrease in absorbance. Experiments were performed in triplicate for each concentration of protein; each line represents the mean, plotted as percent of initial absorbance. (b) Schematic for a PAGE assay to assess if LytH-ActH cleaves crosslinked peptidoglycan; this gel is shown to the right. Uncrosslinked glycan strands (substrate 2) were made from native Lipid II and radiolabeled with [$^{14}$C]-LII$_A^{WTA}$ (substrate 1) at the C6 position (see Supplementary Fig. 8 for additional details)[4]. The radiolabeled glycan strands were crosslinked by *S. aureus* PBP4 to generate crosslinked peptidoglycan (substrate 3)[5]. (c) Peptidoglycan substrates were treated with LytH$^{WT}$-ActH or lysostaphin. Reactions products were resolved by PAGE; this is the full gel for Fig. 3d. Lysostaphin cleaves crosslinks, generating smaller species that migrate faster in the gel.

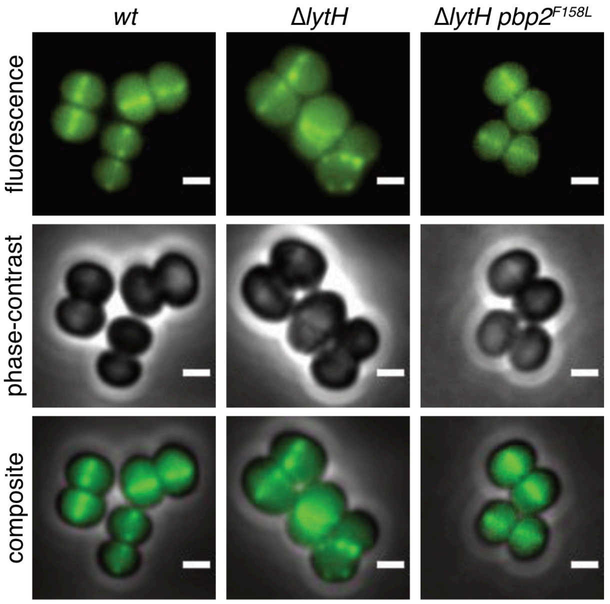


**Supplementary Figure 12:** FtsZ is frequently mislocalized in the absence of LytH. HG003 strains expressing an FtsZ-sGFP sandwich fusion[6] from a cadmium chloride-inducible promoter were grown at 37°C to mid-log phase and imaged using wide-field epifluorescence microscopy. The phase-contrast and composite images for Fig. 4a are provided here. Scale bars, 1 µm.

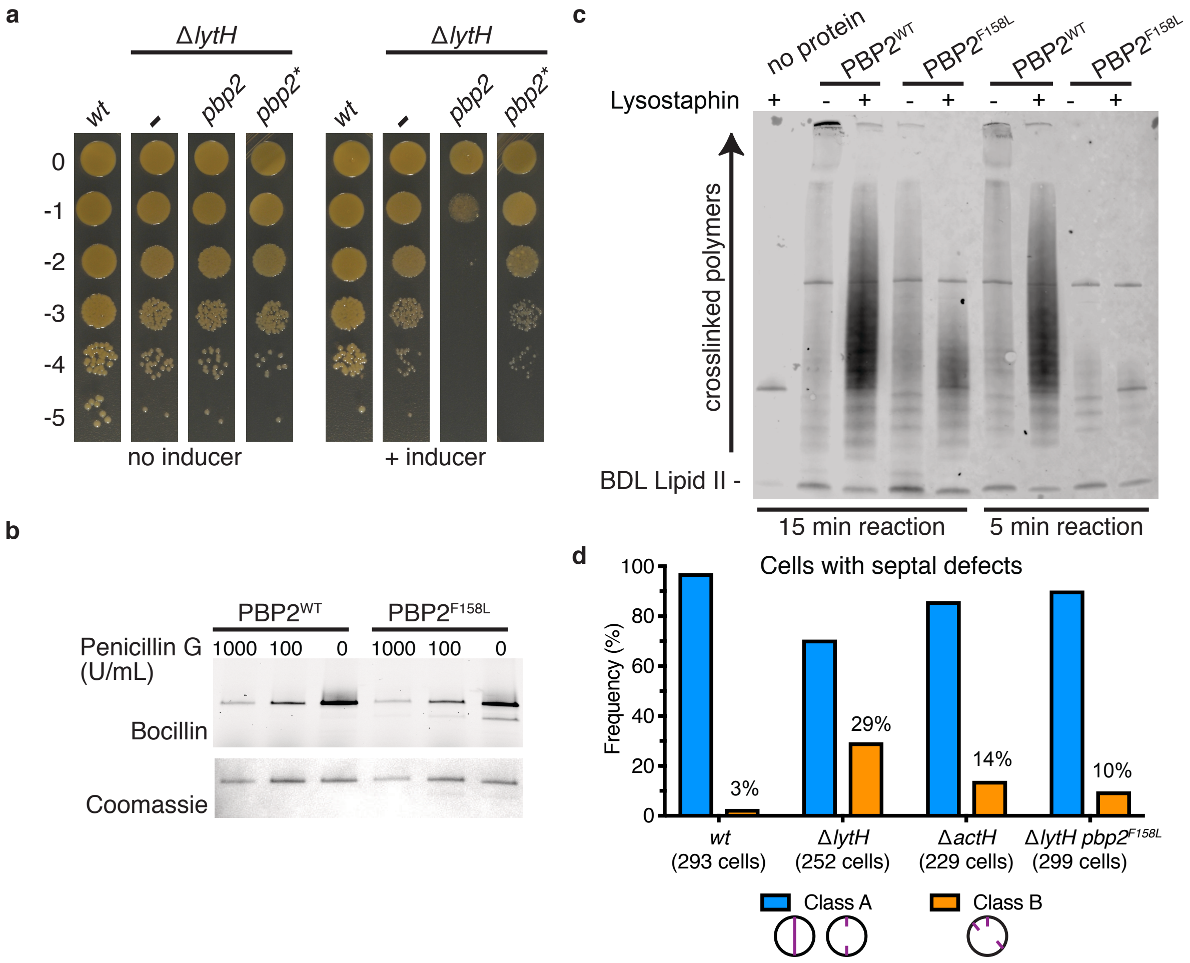


**Supplementary Figure 13:** Reducing PBP2 activity suppresses high-temperature lethality of *lytH* deletion. (a) Spot dilutions showing that excess PBP2 is toxic in the absence of LytH. Wild-type *pbp2* or a *pbp2** suppressor allele (*pbp2*$^{N220\rightarrow KDLN}$) was ectopically expressed from an anhydrotetracycline-inducible promoter in *S. aureus* strain HG003 Δ*lytH* at 37°C. In this strain background, the native *pbp2* WT allele was still present. (b) A representative PBP2 suppressor variant (PBP2$^{F158L}$) was purified. Bocillin-labeling of purified proteins showed that PBP2$^{F158L}$ was properly folded. The Coomassie gel was a control for amounts of protein loaded. (c) A representative western blot comparing PBP2$^{WT}$ and PBP2$^{F158L}$ reactions with Lipid II substrate. PBP2 polymerizes Lipid II into glycan strands and crosslinks the glycan strands; PBP2 also incorporates BDL to enable visualization of crosslinked peptidoglycan. The PBP2 reactions were incubated for 5 or 15 min. When indicated, lysostaphin was added to cleave peptidoglycan crosslinks so that highly-crosslinked material could enter into the gel. This is the full blot for Fig. 4b. (d) A *pbp2* suppressor mutant showed less septal defects. The number of cells with septal defects was counted for the indicated strains.

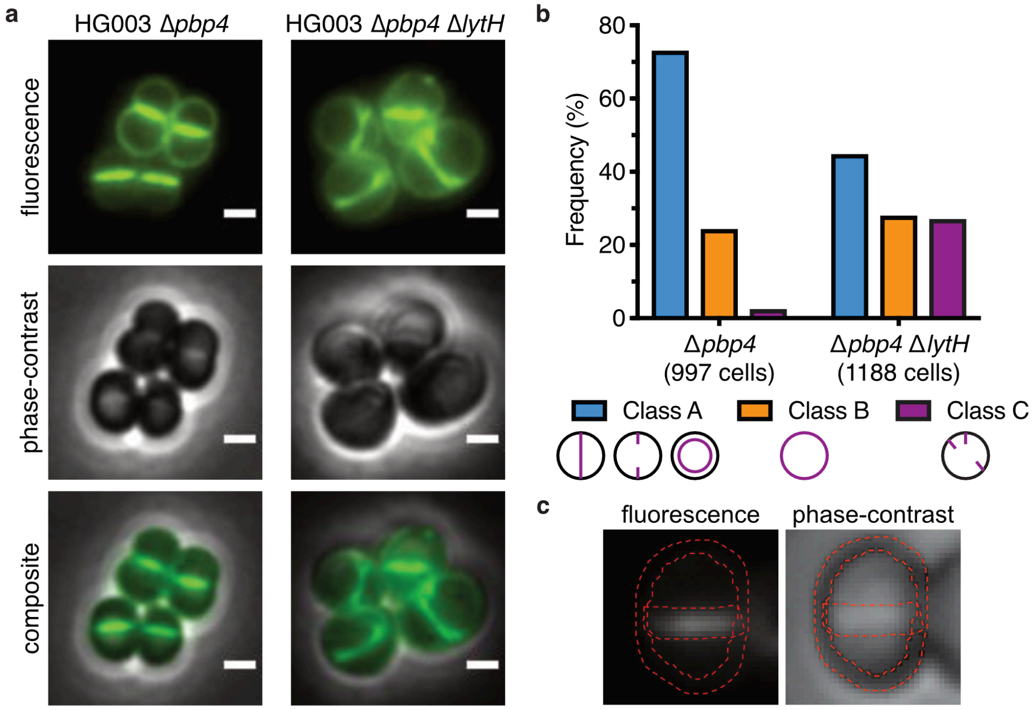


**Supplementary Figure 14:** Loss of LytH leads to accumulation of stem peptides at the cell periphery. (a) Cells grown to mid-log phase at 37°C were labeled with fluorescent D-lysine (FDL) and imaged using wide-field epifluorescence microscopy. FDL is incorporated into peptidoglycan by transpeptidase activity, marking sites of peptidoglycan synthesis[7]. FDL labeling was performed in strains lacking the transpeptidase PBP4 to reduce background[8]. The phase-contrast and composite images for Fig. 5a are provided here. Scale bars, 1 µm. (b) Cells labeled with FDL were quantified and sorted into different classes according to the pattern of FDL staining: at the septum (Class A), around the membrane (Class B), or at misplaced septa and/or punctate foci (Class C). The number of cells counted for each strain are indicated. (c) The ratio of fluorescence intensity at the septum vs. periphery ($F_{septum}/F_{periphery}$) was calculated for cells with a single complete septum, as in Fig. 5a,b and Supplementary Fig. 15. P-values were determined by two-sided Mann-Whitney U tests. To calculate the fluorescence ratio, two regions of interests (ROIs), outlined in red, were defined that encompassed the cell periphery and the septum. These ROIs were super-imposed onto the fluorescence image to retrieve pixel values.

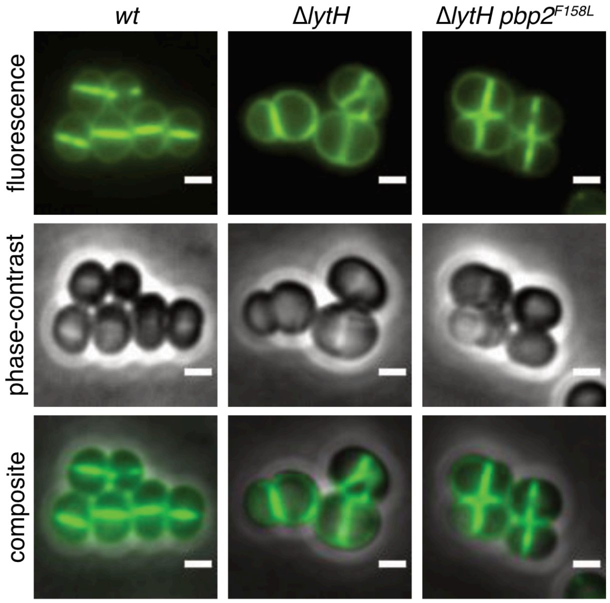


**Supplementary Figure 15:** Loss of LytH leads to increased peptidoglycan synthesis at the cell periphery. HG003 strains were grown to mid-log phase at 37°C in rich media supplemented with excess D-serine, washed, and transferred to rich media without additional D-serine for a short incubation period. Finally, cells were briefly labeled with fluorescent vancomycin and imaged using wide-field epifluorescence microscopy. Because vancomycin has a higher affinity for D-Ala-D-Ala stem peptides than for D-Ala-D-Ser stem peptides[9], which are found in the older cell wall, this assay enables labeling and visualization of newly synthesized peptidoglycan[10]. Fluorescence ratios ($F_{septum}/F_{periphery}$) were calculated for cells with a single complete septum. P-values were determined by two-sided Mann-Whitney U tests. The phase-contrast and composite images for Fig. 5b are provided here. Scale bars, 1 µm.

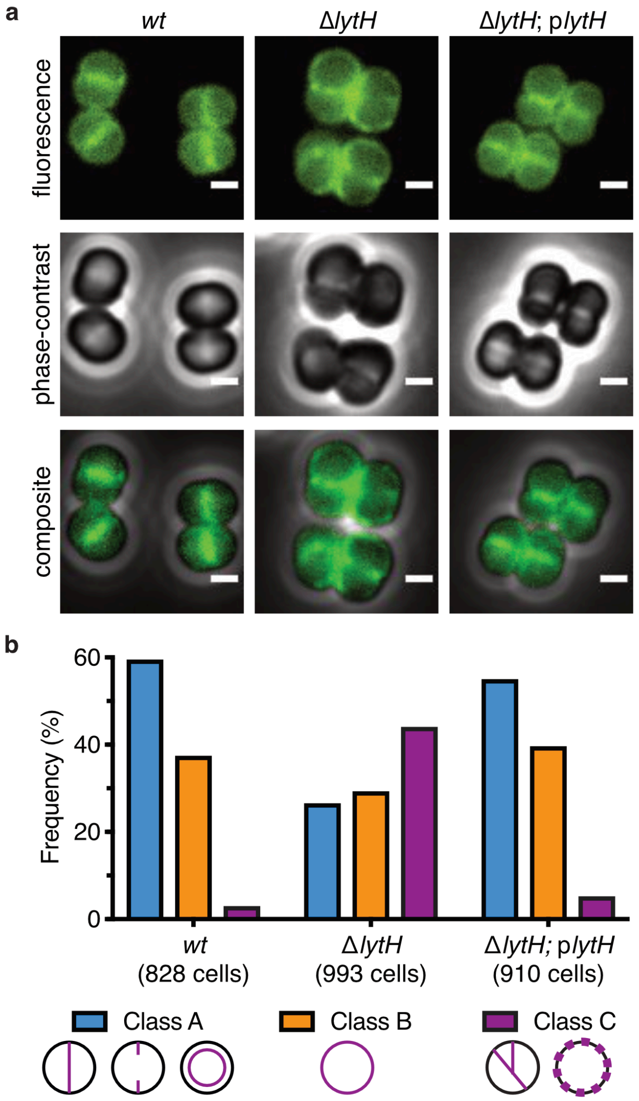


**Supplementary Figure 16:** PBP2 was often mislocalized in the absence of LytH. (a) HG003 strains expressing GFP-PBP2 from a constitutive promoter at the native *pbp2* locus[11] were grown to mid-log phase at 42°C and imaged using wide-field epifluorescence microscopy. To check for complementation, an anhydrotetracycline-inducible $lytH^{WT}$ construct (p*lytH*) was expressed in a Δ*lytH* background. The phase-contrast and composite images for Fig. 5c are provided here. Scale bars, 1 µm. (b) Cells were sorted into different classes according to GFP-PBP2 localization: at the septum (Class A), around the membrane (Class B), or at misplaced septa and/or punctate foci (Class C). The number of cells counted for each strain are indicated.

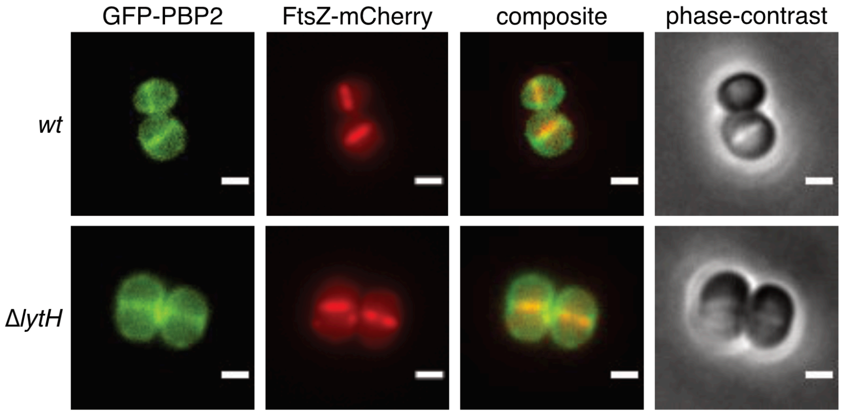


**Supplementary Figure 17:** Colocalization of GFP-PBP2 and FtsZ-mCherry. Dual fluorescent HG003 strains expressing GFP-PBP2[11] from a constitutive promoter and FtsZ-mCherry from an anhydrotetracycline-inducible promoter were imaged by wide-field epifluorescence microscopy. Only cells showing FtsZ signal at the septum were considered for Pearson correlation coefficient (PCC) analysis to calculate the degree of colocalization between GFP-PBP2 and FtsZ-mCherry in a strain, as described in Materials and Methods. P-values were determined by two-sided Mann-Whitney U tests. The two proteins are less colocalized in the absence of LytH. The phase-contrast and composite images for Fig. 5d are provided here. Scale bars, 1 μm.

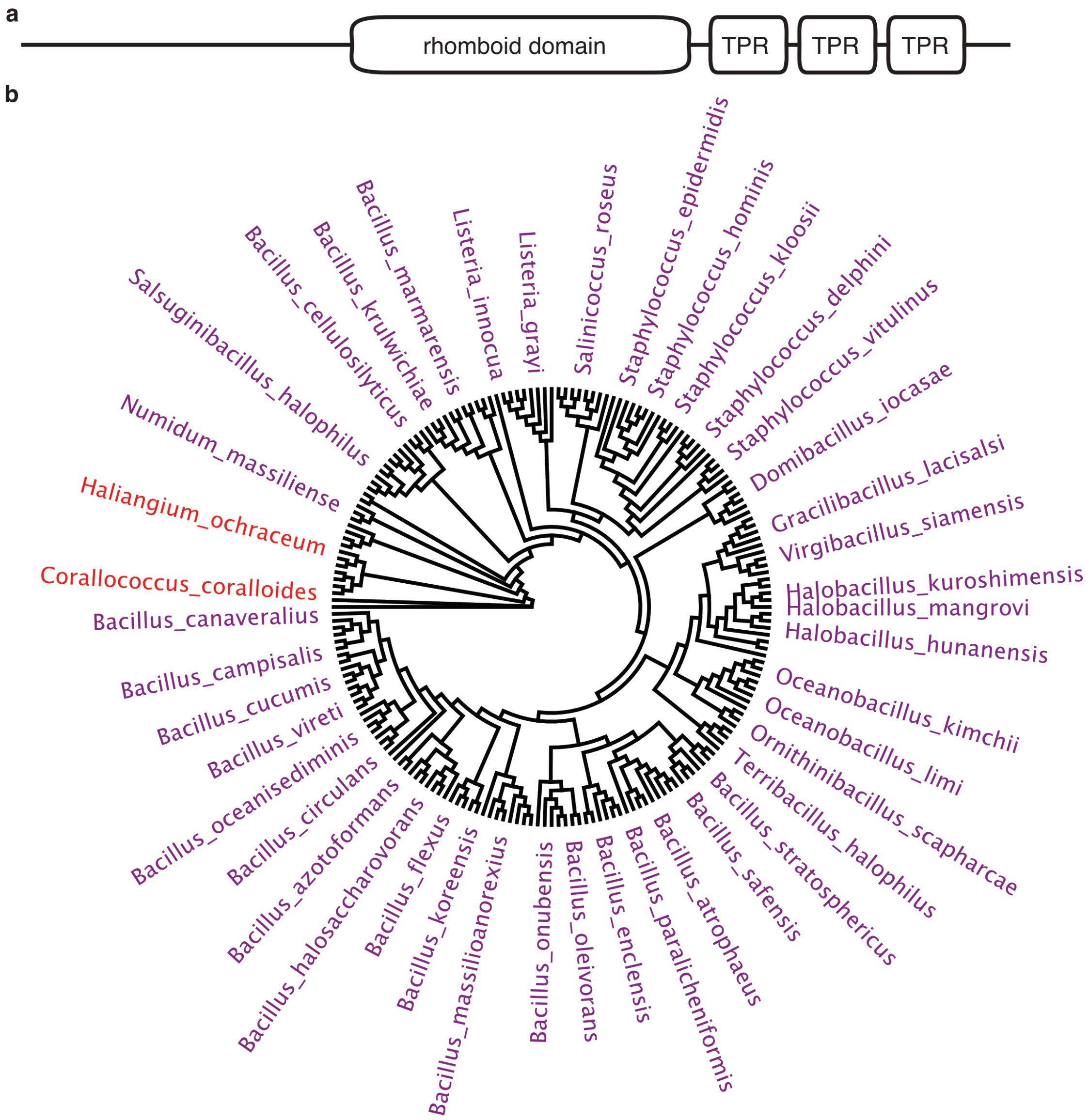


**Supplementary Figure 18:** Homologs of ActH are widespread in Firmicutes. (a) Predicted domain structure of ActH. (b) A cladogram based on 16S rRNA alignment of bacterial species that encode ActH homologs with a similar domain structure was generated. The tree shows 196 bacterial species. All homologs are predicted polytopic membrane proteins bearing a rhomboid domain and tetratricopeptide repeats. For ease of visualization, only some species are labeled. Firmicutes are highlighted in purple and Deltaproteobacteria are highlighted in red.

| **Gene name** | **Annotation**[12] | **Corrected p-value** | **Ratio** |
|---|---|---|---|
| *saouhsc_00023* | Putative protein of unknown function | 2.02E-06 | 0.0564 |
| *saouhsc_00336* | acetyl-CoA acyltransferase | 1.32E-08 | 0.0492 |
| *saouhsc_00445* | Recombination protein RecR | 4.07E-08 | 0.0604 |
| *saouhsc_00499* | Pyridoxal biosynthesis lyase PdxS | 4.49E-12 | 0.0444 |
| *saouhsc_00536* | Branched-chain amino acid aminotransferase IlvE | 2.58E-19 | 0.0218 |
| *saouhsc_00718* | Putative protein of unknown function | 9.99E-07 | 0.0957 |
| *saouhsc_00760* | Diguanylate cyclase GdpS | 3.21E-14 | 0.0469 |
| *saouhsc_00892* | Putative protein of unknown function | 2.20E-05 | 0.0604 |
| *saouhsc_00953* | Diacylglycerol glucosyltransferase UgtP | 8.58E-10 | 0.0411 |
| *saouhsc_01025* | Putative protein of unknown function | 9.54E-16 | 0.0987 |
| *saouhsc_01043* | Dihydrolipoamide dehydrogenase PdhD | 1.84E-07 | 0.0675 |
| *saouhsc_01050* | Putative protein of unknown function | 1.69E-14 | 0.0603 |
| *saouhsc_01154* | Cell division protein SepF | 4.61E-04 | 0.0669 |
| *saouhsc_01612* | 2-oxoisovalerate dehydrogenase, subunit beta | 2.09E-06 | 0.092 |
| *saouhsc_01613* | 2-oxoisovalerate dehydrogenase, subunit alpha | 2.66E-08 | 0.0618 |
| *saouhsc_01649* | Rhomboid-like protein ActH | 2.58E-19 | 0.047 |
| *saouhsc_01739* | Peptidoglycan amidase LytH | 2.30E-10 | 0.0295 |
| *saouhsc_01741* | D-tyrosyl-tRNA deacylase DtdD | 9.31E-05 | 0.0511 |
| *saouhsc_01803* | Putative protein of unknown function | 1.75E-30 | 0.0062 |
| *saouhsc_01827* | Septation ring formation regulator EzrA | 1.11E-10 | 0.0418 |
| *saouhsc_01908* | Putative protein of unknown function | 1.87E-06 | 0.0383 |
| *saouhsc_01960* | Protoporphyrinogen oxidase HemY | 8.51E-04 | 0.0132 |
| *saouhsc_02004* | Putative protein of unknown function | 1.54E-10 | 0.0137 |
| *saouhsc_02049* | PBSX family phage terminase large subunit | 6.14E-03 | 0.0778 |
| *saouhsc_02305* | Alaine racemase Alr | 3.44E-22 | 0.0144 |
| *saouhsc_02337* | UDP-N-acetylglucosamine 1-carboxyvinyltransferase | 5.59E-18 | 0.0345 |
| *saouhsc_02481* | Cobalt transport protein EcfT | 3.25E-05 | 0.0921 |
| *saouhsc_02483* | Cobalt transporter ATP-binding subunit CbiO | 1.27E-06 | 0.0478 |
| *saouhsc_02552* | Putative protein of unknown function | 2.07E-07 | 0.0439 |
| *saouhsc_02571* | Secretory antigen SsaA | 1.20E-10 | 0.0853 |
| *saouhsc_02801* | UTP-glucose-1-phosphate uridylyltransferase GtaB | 1.02E-03 | 0.0717 |
| *saouhsc_02860* | HMG-CoA synthase MvaS | 1.81E-06 | 0.0514 |
| *saouhsc_03049* | Nucleoid occlusion protein Noc | 1.73E-08 | 0.0196 |

**Supplementary Table 1:** A transposon library prepared in the *S. aureus* HG003 strain background was treated with sublethal oxacillin[13,14]. The 10-fold significantly depleted genes represented as green and purple dots in Figure 1b are listed here.

| Unique | Total | Reference | Gene symbol | MW (kDa) | Intensity % |
|---|---|---|---|---|---|
| 10 | 10 | Q2FXU3_LYTH_STAA8 | *lytH* | 32.67 | 11.88 |
| 7 | 10 | Q2FY25_Q2FY25_STAA8 | *saouhsc_01648** | 17.37 | 59.47 |
| 7 | 7 | Q2FY24_Q2FY24_STAA8 | *saouhsc_01649** | 39.37 | 10.99 |
| 3 | 3 | Q2FXZ9_Y1676_STAA8 | *saouhsc_01676* | 35.16 | 1.54 |
| 3 | 3 | Q2G0N0_EFTU_STAA8 | *tuf* | 43.08 | 3.42 |
| 3 | 3 | P60430_RL2_STAA8 | *rplB* | 30.14 | 1.38 |
| 3 | 3 | Q2G247_Y1855_STAA8 | *saouhsc_01855* | 17.99 | 0.70 |
| 3 | 3 | Q2G2D8_Q2G2D8_STAA8 | *saouhsc_00634* | 35.05 | 1.52 |
| 3 | 3 | Q2FW23_RS5_STAA8 | *rpsE* | 17.73 | 0.41 |
| 3 | 3 | Q2G2A5_Q2G2A5_STAA8 | *saouhsc_01041* | 35.22 | 0.54 |
| 2 | 2 | Q2FZ09_Q2FZ09_STAA8 | *recA* | 34.86 | 0.21 |
| 2 | 2 | Q2G0R0_Q2G0R0_STAA8 | *ftsH* | 77.76 | 0.23 |
| 2 | 2 | Q2G0P0_RL1_STAA8 | *rplA* | 24.69 | 0.39 |
| 2 | 2 | Q2FZK0_QOX1_STAA8 | *qoxB* | 75.19 | 0.67 |
| 2 | 2 | Q2FWW9_Q2FWW9_STAA8 | *saouhsc_02152* | 32.93 | 0.23 |
| 2 | 2 | Q2FZV7_Y878_STAA8 | *saouhsc_00878* | 44.08 | 0.13 |
| 2 | 2 | Q2FXT7_Q2FXT7_STAA8 | *saouhsc_01747* | 9.67 | 0.22 |
| 2 | 2 | Q2FZG4_Q2FZG4_STAA8 | *saouhsc_01040* | 41.36 | 1.33 |
| 2 | 2 | Q2FW32_RPOA_STAA8 | *rpoA* | 34.99 | 0.20 |
| 2 | 2 | Q2FW18_RL5_STAA8 | *rplE* | 20.25 | 0.34 |
| 2 | 2 | Q2FW39_RS9_STAA8 | *rpsI* | 14.82 | 0.48 |
| 2 | 2 | Q2FZJ9_QOX2_STAA8 | *qoxA* | 41.75 | 0.29 |
| 2 | 2 | Q2G0V0_Q2G0V0_STAA8 | *saouhsc_00426* | 30.44 | 0.33 |
| 2 | 2 | Q2FVW9_Q2FVW9_STAA8 | *saouhsc_02554* | 33.99 | 0.15 |
| 1 | 2 | Q2G0N9_RL10_STAA8 | *rplJ* | 17.7 | 0.27 |
| 1 | 1 | Q2FWZ8_FTN_STAA8 | *ftnA* | 19.58 | 0.06 |
| 1 | 1 | P0A0G2_RL30_STAA8 | *rpmD* | 6.55 | 0.09 |
| 1 | 1 | Q2G032_Q2G032_STAA8 | *saouhsc_00795* | 36.26 | 0.38 |
| 1 | 1 | Q2FXM9_KPYK_STAA8 | *pyk* | 63.06 | 0.13 |
| 1 | 1 | Q2FYR2_FEMA_STAA8 | *femA* | 49.09 | 0.22 |
| 1 | 1 | Q2FXQ1_RL20_STAA8 | *rplT* | 13.68 | 0.08 |
| 1 | 1 | Q2FXP9_IF3_STAA8 | *infC* | 20.2 | 0.23 |
| 1 | 1 | Q2FXS8_RL21_STAA8 | *rplU* | 11.33 | 0.06 |
| 1 | 1 | Q2FZ25_RS2_STAA8 | *rpsB* | 29.08 | 0.12 |
| 1 | 1 | Q2G0S0_Q2G0S0_STAA8 | *rplY* | 23.77 | 0.20 |
| 1 | 1 | Q2G0Z4_Q2G0Z4_STAA8 | *saouhsc_00367* | 49.38 | 0.34 |
| 1 | 1 | Q2G1G5_PTXBC_STAA8 | *saouhsc_00158* | 50.63 | 0.12 |
| 1 | 1 | Q2G0Q8_Q2G0Q8_STAA8 | *saouhsc_00488* | 32.96 | 0.05 |
| 1 | 1 | Q2FWE9_Q2FWE9_STAA8 | *atpG* | 29.46 | 0.08 |
| 1 | 1 | Q2FYG9_AROC_STAA8 | *aroC* | 43.02 | 0.10 |
| 1 | 1 | Q2G1N4_Q2G1N4_STAA8 | *saouhsc_00074* | 36.72 | 0.12 |
| 1 | 1 | Q2FYT0_Q2FYT0_STAA8 | *saouhsc_01346* | 60.43 | 0.08 |
| 1 | 1 | Q2FXV9_SYA_STAA8 | *alaS* | 98.46 | 0.13 |
| 1 | 1 | Q2FW21_RL6_STAA8 | *rplF* | 19.77 | 0.09 |

**Supplementary Table 2:** Mass spectrometry identifies ActH as a LytH-interaction partner. All peptides identified in the band (outlined by a red box in Fig. 2a) that co-immunoprecipitated with LytH are listed. The top hits (*), *saouhsc_01648* and *saouhsc_01649*, correspond to a single gene, (*actH*). A stop codon was misannotated between the two sequences in the NCTC8325 genome.

| Locus tag | Annotation | Mapped reads | Change | Amino acid change |
|---|---|---|---|---|
| ***Strain TD058*** | | | | |
| | | 453,780 | CAGA -> AGTC | |
| | | 757,505 | G -> T | |
| *saouhsc_01467* | PBP2 | 1,422,211 | (AAAGATTTA)2 -> (AAAGATTTA)3 | N -> KDLN |
| | | 2,350,014 | TCA -> C | |
| ***Strain TD083*** | | | | |
| | | 453,720 | TC -> AT | |
| *saouhsc_00776* | Excinuclease ABC subunit B | 757,486 | GTGCGAAC -> ATTGATGA | VRT -> IDE |
| *saouhsc_00776* | Excinuclease ABC subunit B | 757,494 | G -> C | S -> T |
| | | 758,630 | AC -> GT | |
| | | 758,633 | GAGTCAGCCGAGAAGA -> CGACACCATATACTTT | |
| | | 758,652 | TCCCCG -> GGATTA | |
| *saouhsc_01467* | PBP2 | 1,421,877 | C -> A | A -> E |
| ***Strain TD084*** | | | | |
| *saouhsc_00776* | Excinuclease ABC subunit B | 757,486 | GTGCGAAC -> ATTGATGA | VRT -> IDE |
| *saouhsc_00776* | Excinuclease ABC subunit B | 757,493 | A -> T | S -> C |
| | | 758,652 | TCCCCG -> GGATTA | |
| *saouhsc_01467* | PBP2 | 1,423,062 | C -> T | S -> L |
| ***Strain TD090*** | | | | |
| | | 453,781 | AGA -> GTC | |
| | | 757,505 | G -> T | |
| *saouhsc_01467* | PBP2 | 1,421,859 | C -> T | P -> L |
| ***Strain TD100*** | | | | |
| *saouhsc_01467* | PBP2 | 1,421,945 | G -> A | A -> T |
| | | 2,350,006 | TACTAGA -> ATAG | |
| *saouhsc_02556* | | 2,350,014 | TCA -> C | |
| ***Strain TD101*** | | | | |
| *saouhsc_00776* | Excinuclease ABC subunit B | 757,499 | CCT -> ATG | |
| | | 2,350,006 | TACTAGAC -> ATA | |
| *saouhsc_02556* | Hypothetical protein | 2,350,015 | CA -> TC | |
| *saouhsc_01467* | PBP2 | 1,422,288 | C -> T | A -> V |
| ***Strain TD104*** | | | | |
| *saouhsc_01467* | PBP2 | 1,422,023 | T -> C | F -> L |

**Supplementary Table 3:** Mutations identified by whole-genome sequencing of Δ*lytH* suppressor mutants. The mutations identified in non-coding and protein-coding sequences that showed a variant frequency ≥ 95% are indicated for each suppressor strain.

| **Polymerase** | SgtB | $PBP2^{S398G}$ | $PBP2^{WT}$ |
|---|---|---|---|
| **Transpeptidase** | $PBP2^{E114Q}$ | $PBP2^{E114Q}$ | $PBP2^{WT}$ |
| **% crosslinked species observed (dimer + trimer)** | 29 % | 28 % | 22 % |

**Supplementary Table 4:** PBP2 crosslinks pre-assembled glycan strands. To test if PBP2 can crosslink pre-assembled, linear glycan strands, Lipid II was polymerized with either SgtB or the polymerase-active/transpeptidase-inactive PBP2 variant, $PBP2^{S398G}$. After heat-inactivation of the polymerases, a polymerase-inactive/transpeptidase-active PBP2 variant, $PBP2^{E114Q}$, was added to the reaction mixture. For comparison, Lipid II was also incubated with wild-type PBP2 that both polymerizes and crosslinks. Peptidoglycan polymers were prepared as described in the Materials and Methods, dried, and resuspended in deionized $H_2O$ for LC-MS analysis. Extracted ion chromatograms were generated for the monomer, hydrolyzed monomer, dimer, and trimer. The percentage of crosslinks was determined using the standard equation: sum of (dimer abundance/2 + trimer abundance/3) divided by the total abundance of all muropeptide species observed.

| Gene | Annotated function |
|---|---|
| *saouhsc_01742 (relA)* | (p)ppGpp synthase/hydrolase involved in stringent response |
| *saouhsc_01741 (dtd)* | D-tyrosyl-tRNA deacylase |
| *saouhsc_01739 (lytH)* | Amidase that removes stem peptides from peptidoglycan |
| *saouhsc_01650* | 5-formyltetrahydrofolate cyclo-ligase |
| *saouhsc_01649 (actH)* | Rhomboid-like protein that is an activator of LytH amidase activity |
| *saouhsc_01647* | Putative protein of unknown function |
| *saouhsc_01646 (glk)* | Glucokinase |
| *saouhsc_01645* | Putative protein of unknown function |
| *saouhsc_01644* | Putative protein of unknown function |
| *saouhsc_01643* | Secretion system protein |

**Supplementary Table 5:** *lytH* and *actH* are found in separate operons with genes important for metabolism. Genes that are predicted to be within the same operon as *lytH* or *actH* are indicated with their annotated function[12]. Genes are numbered according to NCTC 8325 nomenclature.

| | Asp | Asn | Ser | Gln | Thr | Gly | Glu | His | Ala | Arg | Tyr | Pro | Met | Val | Trp | Phe | Ile | Lys | Leu |
|---|---|---|---|---|---|---|---|---|---|---|---|---|---|---|---|---|---|---|---|
| **1** | 0.605 | 0.264 | 0.525 | 0.157 | 0.382 | 1.142 | 0.225 | 0.354 | 0.541 | 0.440 | 0.540 | 1.630 | 0.603 | 0.333 | 0.493 | 0.455 | 0.467 | 0.628 | 0.381 |
| **2** | 0.137 | 0.000 | 0.119 | 0.271 | 0.174 | 0.753 | 0.142 | 0.224 | 0.263 | 0.309 | 0.364 | 0.222 | 0.163 | 0.365 | 0.189 | 0.262 | 0.438 | 0.958 | 0.592 |
| **3** | 0.154 | 0.213 | 0.357 | 0.000 | 0.408 | 0.762 | 0.258 | 0.268 | 0.317 | 0.456 | 0.256 | 0.358 | 0.265 | 0.414 | 0.226 | 0.223 | 0.321 | 0.797 | 0.161 |
| **4** | 0.098 | 0.000 | 0.228 | 0.166 | 0.330 | 0.730 | 0.259 | 0.263 | 0.309 | 0.549 | 0.373 | 0.378 | 0.243 | 0.369 | 0.220 | 0.269 | 0.949 | 0.368 | 0.341 |
| **5** | 0.288 | 0.266 | 0.371 | 0.000 | 0.310 | 0.695 | 0.789 | 0.191 | 0.371 | 0.443 | 0.217 | 0.000 | 0.281 | 0.301 | 0.126 | 0.269 | 0.522 | 0.368 | 0.424 |
| **6** | 0.235 | 0.198 | 0.000 | 0.171 | 0.381 | 0.590 | 0.501 | 0.130 | 0.801 | 0.326 | 0.323 | 0.334 | 0.261 | 0.505 | 0.183 | 0.301 | 0.180 | 0.125 | 0.394 |

**Supplementary Table 6:** Edman degradation confirms that LytH is expressed as a full-length protein. The protein band corresponding to C-terminal FLAG-tagged LytH (marked by a purple asterisk in lane 4 from the left in Supplementary Figure 5) was excised for N-terminal sequencing by Edman degradation. Calls for the first six amino acids are listed below as raw Pmol values. The first six amino acids were determined to be: Met-Lys-Lys-Ile-Glu-Ala. Comparing these calls with the published LytH sequence from the HG003 genome[15] confirms that LytH is expressed as a full-length protein with an intact N-terminal transmembrane domain in *S. aureus*.

### Supplementary Table 7: Strains used in this study.

| Strain | Genotype* | Source |
|---|---|---|
| *Escherichia coli* | | |
| XL1-Blue | *recA1 endA1 gyrA96 thi-1 hsdR17 supE44 relA1 lac* [F' *proAB* $lacI^qZo\Delta M15$ Tn*10* ($Tet^R$)] | Agilent Technologies |
| NEB 10-beta | *Δ(ara-leu) 7697 araD139 fhuA ΔlacX74 galK16 galE15 e14-* *ϕ80*d*lacZΔM15 recA1 relA1 endA1 nupG rpsL* ($Str^R$) *rph spoT1* Δ(*mrr-hsdRMS-mcrBC)* | New England Biolabs |
| Stellar | *F- endA1 supE44 thi-1 recA1 relA1 gyrA96 phoA ϕ80d lacZΔ M15 Δ(lacZYA-argF) U169 Δ(mrr-hsdRMS-mcrBC) ΔmcrA λ-* | Takara Clontech |
| BL21(DE3) | *F- ompT* $hsdS_B$($r_B$- $m_B$-) *gal dcm* (DE3) | Novagen |
| C43(DE3) | *F- ompT* $hsdS_B(r_B\text{- } m_B\text{-})$ *gal dcm* (DE3) | Lucigen |
| DC10B | *F- mcrA Δ(mrr-hsdRMS-mcrBC) ϕ80*d*lacZΔ*M15 *ΔlacX74 endA1 recA1 Δ(ara-leu)7697 araD139 galU galK nupG rpsL λ- Δdcm* | [16] |
| NovaBlue(DE3) | *endA1 hsdR17* ($r_{k12}$- $m_{k12}$+) *supE44thi-1 recA1 gyrA96 relA1 lac* (DE3) *F' proAB* $lacI^QZ\Delta M15$*::Tn10* ($Tet^R$) *gyrA96*($nal^R$) *thi-1 recA1 relA1 lac glnV44 F'*[*::Tn10* $proAB^+$ $lacI^q$ Δ(*lacZ*)*M15*] *hsdR17*($r_K$– $m_K$+) | Novagen |
| *Staphylococcus aureus* | | |
| RN4220 | Wild-type | [17] |
| HG003 | Wild-type | [18] |
| TD011 | RN4220 (pTP44), $tet^R$ | [19] |
| TD024 | HG003 Δ*lytH*::$kan^R$ | This study |
| TD036 | HG003 Δ*lytH*::$kan^R$ (pTD10) | This study |
| TD037 | HG003 Δ*lytH*::$kan^R$ (pTD11) | This study |
| TD058 | HG003 Δ*lytH*::$kan^R$ $pbp2^{N220\rightarrow KDLN}$ suppressor | This study |
| TD074 | HG003 Δ*lytH*::$kan^R$ (pTD18) | This study |
| TD075 | HG003 Δ*lytH*::$kan^R$ (pTD19) | This study |
| TD083 | HG003 Δ*lytH*::$kan^R$ $pbp2^{A109E}$ suppressor | This study |
| TD084 | HG003 Δ*lytH*::$kan^R$ $pbp2^{S504L}$ suppressor | This study |
| TD090 | HG003 Δ*lytH*::$kan^R$ $pbp2^{P103L}$ suppressor | This study |
| TD100 | HG003 Δ*lytH*::$kan^R$ $pbp2^{A132T}$ suppressor | This study |
| TD101 | HG003 Δ*lytH*::$kan^R$ $pbp2^{A246V}$ suppressor | This study |
| TD104 | HG003 Δ*lytH*::$kan^R$ $pbp2^{F158L}$ suppressor | This study |
| TD134 | HG003 Δ*lytH*::$kan^R$ (pTD30) | This study |
| TD135 | HG003 Δ*lytH*::$kan^R$ (pTD31) | This study |
| TD140 | RN4220 Δ*tarO*, unmarked deletion | [20] |
| TD157 | HG003 Δ*lytH*::$kan^R$ (pTD34) | This study |
| TD164 | HG003 Δ*lytH*::$kan^R$ (pTD39) | This study |
| TD177 | HG003 Δ*saouhsc_01649*::$tet^R$ | This study |
| TD178 | HG003 Δ*lytH*::$kan^R$ Δ*saouhsc_01649*::$tet^R$ | This study |
| TD179 | HG003 Δ*lytH*::$kan^R$ Δ*saouhsc_01649*::$tet^R$ (pTD30) | This study |
| TD215 | HG003 Δ*pbp4*, unmarked deletion | Ting Pang |
| TD240 | HG003 Δ*pbp4* Δ*lytH*::$kan^R$ | This study |
| TD261 | HG003 (pPBP2-31) | This study |
| TD262 | HG003 Δ*lytH*::$kan^R$ (pPBP2-31) | This study |
| TD263 | HG003 Δ*lytH*::$kan^R$ (pTD10; pPBP2-31) | This study |
| TD268 | HG003 (pTD71) | This study |
| TD269 | HG003 Δ*lytH*::$kan^R$ (pTD71) | This study |
| TD278 | HG003 (pPBP2-31; pTD72) | This study |
| TD279 | HG003 Δ*lytH*::$kan^R$ (pPBP2-31; pTD72) | This study |
| TD282 | HG003 Δ*lytH*::$kan^R$ $pbp2^{F158L}$ (pTD71) | This study |

*See Supplementary Table 8 for list of abbreviations.

**Supplementary Table 8: Plasmids used in this study.**

| Plasmid | Description* | Source |
|---|---|---|
| pTP63 | atc-inducible, integrative *cam*$^R$ expression vector for *S. aureus* | [19] |
| pLOW | IPTG-inducible, low-copy *erm*$^R$ expression vector for *S. aureus* | [21] |
| pKFC | *cam*$^R$ vector for making gene deletions in *S. aureus* | [22] |
| pTM186 | *cam*$^R$ vector for amplification of GFPmut2 sequence | Timothy C. Meredith |
| pET15b | *amp*$^R$ protein expression vector | Novagen |
| pET42a(+)-*pbp2* | *kan*$^R$ protein expression vector for *S. aureus* PBP2 [K60-S716] | [23] |
| pET42a(+)-*pbp2*$^{S398G}$ | *kan*$^R$ protein expression vector for *S. aureus* PBP2$^{S398G}$ [K60-S716] | [23] |
| pET42a(+)-*pbp2*$^{E114Q}$ | *kan*$^R$ protein expression vector for *S. aureus* PBP2$^{E114Q}$ [K60-S716] | [24] |
| pTarKO | *cam*$^R$ vector for one-step gene deletions in *S. aureus* | [20] |
| pET28b(+) | *kan*$^R$ protein expression vector | Novagen |
| pKFC-*tarO* | *cam*$^R$ plasmid for constitutive expression of *tarO* in *S. aureus* | [20] |
| pETDuet-1 | *amp*$^R$ vector for dual protein expression | EMD Millipore |
| pMgt1 | *amp*$^R$-marked *S. aureus* SgtB$^{WT}$-His$_6$ expression vector | [2] |
| pET24b(+)-*sgtB*$^{Y181D}$ | *amp*$^R$-marked *S. aureus* SgtB$^{Y181D}$-His$_6$ expression vector | [2] |
| pLcpB | *kan*$^R$-marked *S. aureus* His$_8$-LcpB [S31-N405] expression vector | [4] |
| pMW1010 | pET28b(+)-His$_6$-*EF_3129* [T36-P429] expression vector for *E. faecalis* PBPX, *kan*$^R$ | [3] |
| pCN51 | $CdCl_2$-inducible, replicative *erm*$^R$ expression vector for *S. aureus* | [25] |
| pCN-ftsZ$^{55\text{-}56}$sGFP | $CdCl_2$-inducible vector to express FtsZ-sGFP sandwich fusion | [6] |
| pPBP2-31 | *erm*$^R$ vector for constitutive expression of GFP-PBP2 | [11] |
| pTD2 | pET15b-His$_6$-*lytH* [E41-A291] expression vector | This study |
| pTD3 | pET15b-His$_6$-*lytH* [T102-A291] expression vector | This study |
| pTD6 | pKFC vector to make Δ*lytH*::*kan*$^R$ marked deletion in *S. aureus* | This study |
| pTD10 | pTP63-*lytH* [M1-A291]-1x FLAG containing *lytH* RBS | This study |
| pTD11 | pTP63-*gfpmut2* [M1-K238]-1x FLAG containing *lytH* RBS | This study |
| pTD18 | pTP63-*pbp2* [M1-N727]-1x FLAG containing *pbp2* RBS | This study |
| pTD19 | pTP63-*pbp2*$^{N220\rightarrow KDLN}$ [M1-N730]-1x FLAG containing *pbp2* RBS | This study |
| pTD30 | pLOW-*lytH* [M1-A291]-1x FLAG containing *rpoB* RBS | This study |
| pTD31 | pLOW-*gfpmut2* [M1-K238]-1x FLAG containing *rpoB* RBS | This study |
| pTD34 | pLOW-*lytH* [M1-K120]-1x FLAG containing *rpoB* RBS | This study |
| pTD39 | pLOW-*lytH*$^{D195A}$ [M1-A291]-1x FLAG containing *rpoB* RBS | This study |
| pTD42 | pET28b(+)-*lytH* [M1-A291]-His$_6$ expression vector | This study |
| pTD47 | pTarKO vector to make Δ*saouhsc_01649*::*tet*$^R$ marked deletion | This study |
| pTD48 | pET42a(+)-*pbp2*$^{F158L}$ [K60-S716]-His$_8$ expression vector | This study |
| pTD51 | pETDuet-1-His$_6$-*saouhsc_01649* [N2-K487]-*lytH* [M1-A291]-1x FLAG dual expression vector | This study |
| pTD52 | pET28b(+)-His$_6$-*saouhsc_01649* [N2-K487] expression vector | This study |
| pTD54 | pETDuet-1-His$_6$-*saouhsc_01649* [N2-K487]-*lytH*$^{D195A}$ [M1-A291]-1x FLAG dual expression vector | This study |
| pTD71 | pCN51-*ftsZ*$^{55\text{-}56}$*sGFP* for $CdCl_2$-inducible expression of FtsZ-sGFP sandwich fusion in *S. aureus* | This study |
| pTD72 | pTP63-*ftsZ* [M1-R390]-*mCherry* containing native *ftsZ* ribosome-binding site and 5-amino acid linker (SCGAS) | This study |

*Abbreviations: *amp*$^R$, ampicillin/carbenicillin resistance; *cam*$^R$, chloramphenicol resistance; *kan*$^R$, kanamycin resistance; *erm*$^R$, erythromycin resistance; *tet*$^R$, tetracycline resistance; atc, anhydrotetracycline; IPTG, isopropyl-β-D-1-thiogalactopyranoside; RBS, ribosome-binding site

**Supplementary Table 9: Oligos used in this study.**

| Primer | Sequence (5' to 3') |
|---|---|
| oTD3 | gcgaaccatttgaggtgataggtaagattat |
| oTD4 | tcctaggtactaaaacaattcatccagtaa |
| oTD7 | ttactggatgaattgttttagtacctaggaggcttgcaaaaatatgtgaaagtagttatc |
| oTD8 | tgcaggtcgaccactaacgcttcgttatattcttttcg |
| oTD15 | tgccgcgcggcagccatatggaagatagtgggaacatcacgataac |
| oTD16 | tgccgcgcggcagccatatgacaaatttagatattgtcgcggataatacg |
| oTD17 | ctttgttagcagccggatccctacgcagaaaaataaattttaaggccatc |
| oTD24 | cccggggatccctgaaagacaacgtattttagataaacaacg |
| oTD25 | ataatcttacctatcacctcaaatggttcgctcattgaattttgccgtcctgactttcat |
| oTD38 | atgatggtaccatgaaagtcaggacggcaaaattcaatg |
| oTD39 | cggcgctcagcctacttgtcgtcatcgtctttgtagtccgcagaaaaataaattttaaggccatcaac |
| oTD40 | atgatggtaccatgaaagtcaggacggcaaaattcaatgagtaaaggagaagaacttttcactgg |
| oTD41 | cggcgctcagcctacttgtcgtcatcgtctttgtagtctttgtatagttcatccatgccatgtgtaa |
| oTD54 | tatgatggtacctattagatgaaagtgaggaccgcgtatg |
| oTD56 | tttaaatcttttaaatctttattaaagtaatacttagcagcagc |
| oTD57 | agatttaaacttagcggaagaagcttatttagccgg |
| oTD58 | ccggcgctcagcttacttgtcgtcatcgtctttgtagtcgttgaatatacctgttaatccaccgctg |
| oTD85 | cccggggatccctacttgtcgtcatcgtctttgtagtc |
| oTD86 | tgcaggtcgaccataatttttgaggggtgaatctgtatgaaaaaaatagaggcatggttatctaaaaaggg |
| oTD87 | tgcaggtcgaccataatttttgaggggtgaatctgtatgagtaaaggagaagaacttttcactgga |
| oTD103 | attatgtatactcaaataggcatcgcctttgatatc |
| oTD104 | gctgcgttagaatcatctaatgcaaatggaatgacag |
| oTD106 | cccggggatccctacttgtcgtcatcgtctttgtagtccgcagaaaaataaattttaaggccatcaac |
| oTD111 | tccatttgcattagatgattctaacgcagcattatgtatactcaaataggcatcgcctttgatat |
| oTD116 | tggtgctcgaggctgccgcgcggcaccagtgcggccgccgcagaaaaataaattttaaggccatcaac |
| oTD117 | atataccatgggcaaaaaaatagaggcatggttatctaaaaaggg |
| oTD141 | cccggggatccagcgcatggttgtaaaattttaactttgca |
| oTD142 | caactcaaattgcgagatttgggttgttagtcctccactatgctgcttgata |
| oTD143 | gatagataaagtaagatatatgttcaataaaataacttagatgagtcgaaaaataaataacttttatgatgtacaac |
| oTD144 | tgcaggtcgaccctagaacgatatatttcggattacttgta |
| oTD145 | caacccaaatctcgcaatttgagttg |
| oTD146 | ctaagttattttattgaacatatatcttactttatctatc |
| oTD148 | tgcatctttaacaacttgttgtgttaatgttgag |
| oTD149 | cttttatcacaacataaatctattggacgtaaagct |
| oTD160 | atccgaattcgaacatagacaaacaattttggaaaacaatatattattg |
| oTD161 | ccgcaagcttttatttatttttcgactcatttgatttagtcaactc |
| oTD166 | tagttaagtataagaaggagatatacatatgaaaaaaatagaggcatggttatctaaaaagg |
| oTD167 | cagcggtttctttaccagactcgagggtaccctacttgtcgtcatcgtctttgtagtc |
| oTD219 | ggtggtggtagcgctggtggagctatgaaaaaaatagaggcatggttatctaaaaagg |
| oTD220 | ctacgcagaaaaataaattttaaggccatc |
| oTD221 | tgatagagtatgatggtaccatgaaagtcaggacggcaaaattc |
| oTD222 | agcgaccggcgctcagcctacgcagaaaaataaattttaaggcc |
| oTD223 | atgaaagtcaggacggcaaaattcaatggtgagcaagggcgagga |
| oTD224 | agctccaccagcgctaccaccacccttgtacagctcgtccatgcc |
| oTD227 | agcgaccggcgctcagcctacttgtacagctcgtccatgcc |
| oTD248 | tgatagagtatgatggtaccggccaataaaactaggaggaaatttaaatg |
| oTD249 | ggaggcgccgcaggaacgtcttgttcttcttgaacgtctttc |
| oTD250 | acgttcctgcggcgcctccgctattatcaaagaatttatgagattcaaagttc |
| YQ1 | acgcagtacttgcaactcaggacaatcgtttctacgaacatg |
| YQ2 | catgttcgtagaaacgattgtcctgagttgcaagtactgcgt |
